# Perception and propagation of activity through the cortical hierarchy is determined by neural variability

**DOI:** 10.1101/2021.12.28.474343

**Authors:** James M. Rowland, Thijs L. van der Plas, Matthias Loidolt, Robert M. Lees, Joshua Keeling, Jonas Dehning, Thomas Akam, Viola Priesemann, Adam M. Packer

## Abstract

The brains of higher organisms are composed of anatomically and functionally distinct regions performing specialised tasks; but regions do not operate in isolation. Orchestration of complex behaviours requires communication between brain regions, but how neural activity dynamics are organised to facilitate reliable transmission is not well understood. We studied this process directly by generating neural activity that propagates between brain regions and drives behaviour, allowing us to assess how populations of neurons in sensory cortex cooperate to transmit information. We achieved this by imaging two hierarchically organised and densely interconnected regions, the primary and secondary somatosensory cortex (S1 and S2) in mice while performing two-photon photostimulation of S1 neurons and assigning behavioural salience to the photostimulation. We found that the probability of perception is determined not only by the strength of the photostimulation signal, but also by the variability of S1 neural activity. Therefore, maximising the signal-to-noise ratio of the stimulus representation in cortex relative to the noise or variability in cortex is critical to facilitate activity propagation and perception. Further, we show that propagated, behaviourally salient activity elicits balanced, persistent, and generalised activation of the downstream region. Hence, our work adds to existing understanding of cortical function by identifying how population activity is formatted to ensure robust transmission of information, allowing specialised brain regions to communicate and coordinate behaviour.

---

Key to the orchestration of behaviour by neural systems is that information, in the form of neural activity, is reliably and accurately transmitted between anatomically distinct brain regions performing specialised tasks. Activity is transformed at each stage of its journey, and circuits often perform multiple tasks in parallel^1,2^, so it is challenging to disambiguate which facets of neural activity contribute to a specific behaviour or process. In-depth analysis of cellular resolution recordings of neural activity during sensory stimulation^3–7^ and/or well-characterised behaviour^8–12^ has begun to disentangle how sensation, decisions, and actions are encoded in individual brain regions. However, how sensory-related activity is structured to facilitate its journey through the brain, and how it interacts with ongoing cortical activity, is less well understood. The relationship between sensory-related and ongoing cortical activity is crucial to understand as, *in silico*, the structure and variability of ongoing activity, which varies with internal state (Tanashiro et al. 2017, Wilting et al. 2018, Shi et al. 2022), can facilitate or limit the detection or classification of simple stimuli (Ecker et al. 2016, Zylberberg et al. 2018, Cramer et al. 2020, Bernardie et al 2021). *In vivo* experiments have shown that the influence of ongoing cortical activity on sensory and behavioural responses can be of similar importance as stimulus strength^13–16^.

One of the overarching functions of neural systems is to detect and respond to stimuli originating from the outside world. This requires reliable propagation of neural signals from sensory organs through anatomical hierarchies in the brain. Signal propagation is usually studied in psychophysical assays where stimuli of increasing strength are presented to the participant, and their detection signalled behaviourally, while the activity of relevant brain regions is recorded to capture propagating activity^17–23^.

However, a challenge in quantifying signal propagation is that correlated activity in two regions may not reflect causal influence of one on the other, but rather may reflect common input^24–27^. Taking causal control of neural activity by directly stimulating neurons circumvents this issue, as activity locked temporally to stimulation of a connected region likely arises from propagation. This can be achieved by driving spikes directly through stimulation of individual neurons^28–30^ or groups of neurons^31–35^. However, so far, detection of optogenetic stimulation with single-cell precision has only been paired with single-region neural recordings^32,33^, limiting signal propagation studies to within-region dynamics.

Here, we harness the single-cell precision of all-optical interrogation^36^ to investigate how the collective dynamics of neural activity determine activity propagation and guide behaviour. We imaged two hierarchically organised, densely interconnected and functionally well-characterised^37–44^ regions, the primary and secondary somatosensory cortex (S1 and S2) while performing two-photon optogenetic photostimulation of S1 neurons, and trained mice to report stimulation. By recording neural activity occurring before and after photostimulation in both brain regions simultaneously, we were able to assess how behaviourally salient stimulation is propagated through anatomically distinct brain regions and interacts with their ongoing activity.

## Causally driving behaviour with all-optical interrogation across cortical areas

We developed a preparation that allowed us to record activity in two brain regions (S1 and S2) simultaneously, while holographically photostimulating S1 **(Fig. 1a)**. To achieve this, we expressed the genetically encoded calcium indicator GCaMP6s^45^ and the somatically targeted, red-shifted opsin C1V1-Kv2.1^29,46^ in layer 2/3 across both S1 and S2. We localised S1 and S2 by performing wide-field calcium imaging during deflection of individual whiskers **(Fig. 1b, Extended Data Fig. 1)**. A field of view was selected which spanned the cortical representations of multiple whiskers across the a, b and c rows in both S1 and S2. Mice were head-fixed and we utilised a two-photon microscope, based on previous designs^36^, adapted to perform two-photon calcium imaging of neural activity across a large field of view (1.35 mm diameter), while performing two-photon (2P) photostimulation of S1 neurons. Before each experiment, we photostimulated all opsin-expressing neurons in groups of 20 to find cells responsive to photostimulation, with neurons within a 350 μm diameter targeted simultaneously **(Fig. 1c)**. Targeted neurons, defined as cells within 15 µm of the centre of a photostimulation beam, elicited significant excitatory responses while nearby S1 non-target neurons elicited significant inhibitory responses on average (**Fig. 1d, Extended Data Fig. 2a,b**, Wilcoxon signed-rank test, p < 0.05, Bonferroni corrected).

**Figure 1:**
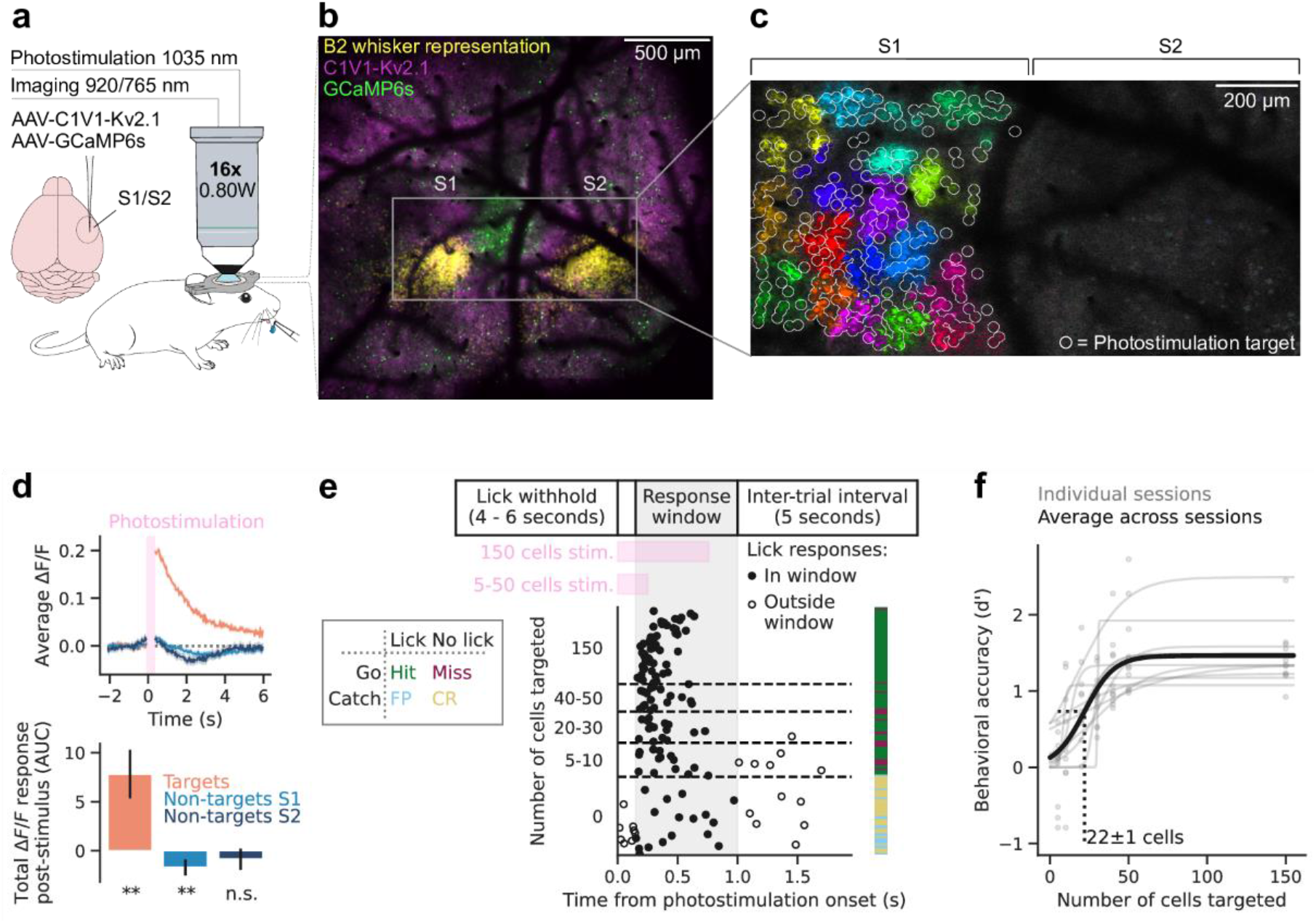
Recording neural activity in S1 and S2 during behavioural report of targeted two-photon photostimulation of S1. **(a)** Schematic^107^ of experimental setup. *Left:* viral strategy for expression of GCaMP6s and C1V1-Kv2.1-mScarlet in S1 and S2. *Right:* mice with a cranial window installed over S1 and S2 were headfixed under a two-photon microscope. A lick spout was placed within reach of the tongue, through which the animal reported perception of photostimulation by licking and received a water reward. **(b)** Example imaging field of view used to localise S1 and S2 by whisker stimulation (stimulus-triggered average widefield calcium whisker response shown in *yellow*, see **Extended Data Fig. 1** for multiple whiskers), overlaid with aligned two-photon images of GCaMP6s (*green*) and C1V1-Kv2.1-mScarlet (*magenta*) expression. **(c)** Example two-photon calcium imaging field of view with photostimulation targets. The intensity of each pixel is proportional to the change in fluorescence intensity post-photostimulation compared to pre-photostimulation (stimulus triggered average, **methods**); bright pixels indicate a photostimulation-induced increase in calcium activity. Pixels are colour-coded based on whether they were photostimulated simultaneously. Non-targeted cells, including those in S2, are not visible because different cells would respond to repeated stimulation of the same group of targeted cells and were therefore averaged out. **(d)** Top: Example activity responses to photostimulation of a single recording session. Orange shows the response to photostimulation of cells directly targeted with light, averaged across cells and across trials. Light and dark blue show the response of cells not directly targeted in S1 and S2 respectively. Only trials in which photostimulation was delivered were included. Data are blanked while the photostimulation laser was on (pink bar) as this causes a large artifact unrelated to neural activity. Bottom: the total ΔF/F activity post-stimulus is shown, as defined by the Area Under the Curve (AUC) of the traces of the top panel, for all 11 recording sessions (mean ± 95% confidence interval across sessions). We tested whether each condition was significantly different from 0 (Wilcoxon signed-rank test, significance is indicated by: ** (p < 0.01) or n.s. (not significant)). **(e)** *Top:* Timing of a single behavioural trial. *Bottom left:* Behavioural response matrix. *Bottom right:* Example lick raster from a single session sorted by number of cells targeted and by time within each bin. Each row of the plot shows the first lick within an individual trial. The colour bar shows the outcome of the trial as defined in the behavioural response matrix. **(f)** Psychometric curves showing behavioural performance (d’) as a function of the number of cells targeted by photostimulation. Each grey point is the d’ computed for a given number of cells targeted for an individual session and each grey line is a logistic function fit for an individual session. The thick blackline shows the fit for all data points across all sessions (N = 11 sessions; N = 5 mice). The grey dashed line shows that the 50% point from the fit across all sessions occurs at 22 ± 1 cells targeted.

We trained mice via operant conditioning to associate direct photostimulation of neurons in S1 with water reward. After initial training, mice reported photostimulation by licking a spout **(Fig. 1e)**. Crucially, to prevent a guessing strategy and/or random licking, mice had to withhold licking to initiate a trial after a variable inter-trial interval. Successful report of stimulation on ‘go’ trials was scored as a ‘hit’ and the mouse was rewarded; trials in which the animal failed to report the stimulation through licking were scored as a ‘miss’. We also randomly interleaved ‘catch’ trials in which no stimulus was delivered to analyse the detection performance relative to chance. Catch trials in which the animal licked were ‘false positive’ trials while trials when the animal appropriately refrained from licking were ‘correct rejection’ trials **(Fig. 1e)**. Between 5 and 150 photoresponsive S1 neurons were targeted on a single go trial. The number and identity of cells targeted were varied randomly on each trial to generate a psychometric curve **(Fig. 1f)**, which shows that stimulating a greater number of neurons increased licking behaviour accuracy. The 50% point of the psychometric curve fit across all sessions is 22 ± 1 neurons (n = 11 sessions recorded from 5 mice, ± 95% confidence interval), in accordance with previously published findings^32^. Thus, we have developed a preparation in which activity that causally drives behaviour can be recorded, with single-cell resolution, both locally and, after propagation downstream.

## 2P photostimulation of S1 drives balanced activity that propagates to S2

To understand how neural activity is propagated between brain regions to guide behaviour, we compared neural responses during hit and miss trials, both in the directly stimulated region (S1) and the downstream region (S2). Qualitatively, hit trials elicit a diverse array of relatively strong excitatory and inhibitory responses both in S1 and S2, whereas both excitatory and inhibitory responses were less strong on miss trials **(Fig. 2a; Extended Data Fig. 3,4)**. Excited and inhibited cells are also observed in ‘reward only’ trials **(methods)** in which reward was delivered in the absence of photostimulation, however, the neuronal responses are less pronounced compared to hit trials, indicating that the response to reported stimulation is not just explained by neural activity driven by reward **(Fig. 2a; Extended Data Fig. 3,4)**.

**Figure 2:**
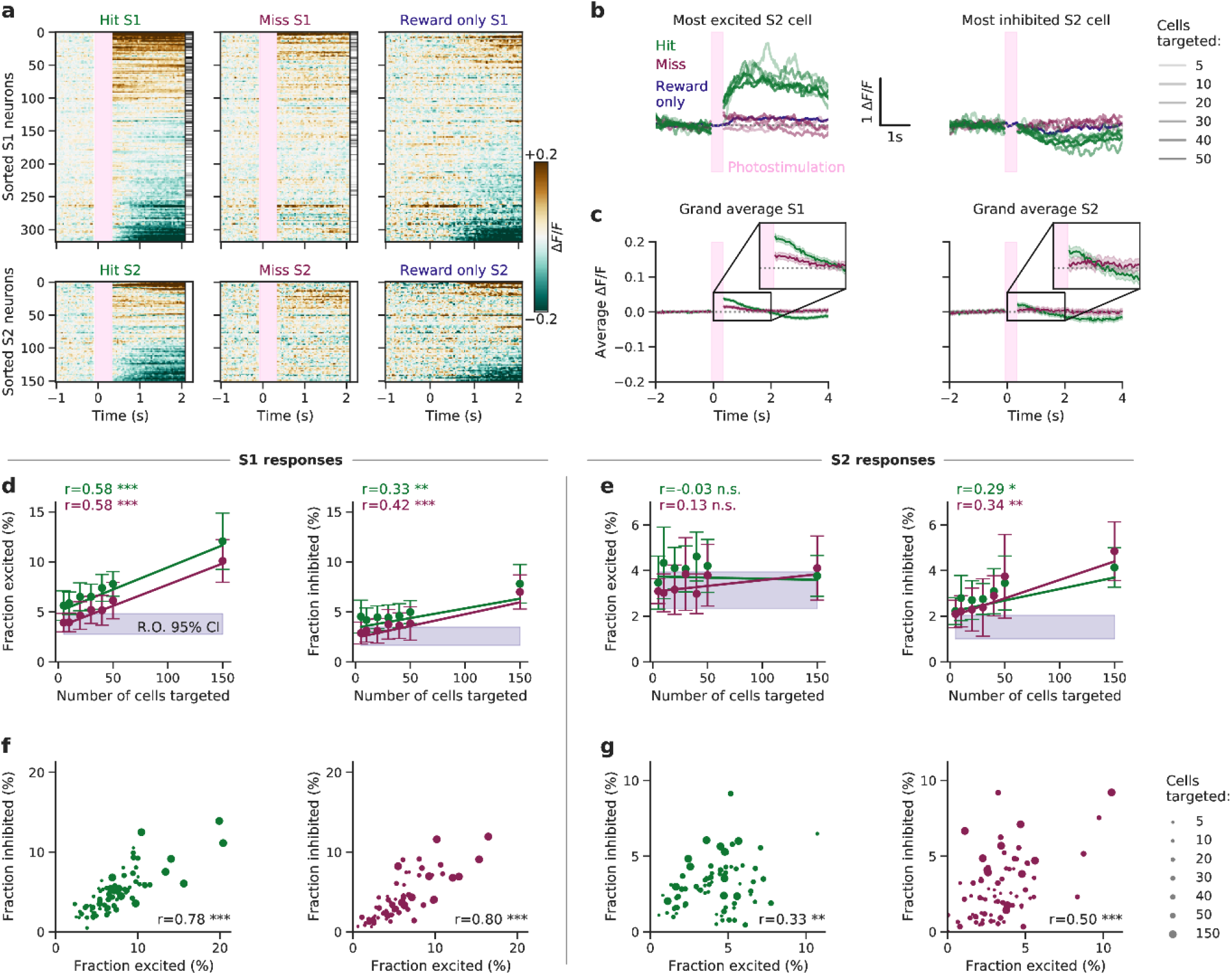
Activity driven by targeted two-photon photostimulation is propagated from S1 to S2. **(a)** Raster plots showing the trial-averaged response of different trial types (*Left:* hit, *Middle*: miss, *Right:* reward only) to photostimulation (pink vertical bar, hit/miss) and/or reward (hit/reward only) of individual cells from a single session. (All trial types of all sessions are shown in **Extended Data Fig. 3,4**). Cells are sorted by the sum of their trial-averaged responses across all three trial types **(methods)**. Clear excitatory and inhibitory responses are elicited in S1 and S2 on hit trials that are not observed on miss trials or reward only trials. The intensity of the grey-scale bar on the right hand side of the hit and miss rasters is proportional to the number of times each cell was directly targeted by the photostimulation beam, for hit and miss trials separately. Trials in which 150 cells were targeted were removed for display because their stimulation period is longer. Data are bound between -0.2 and +0.2 ΔF/F and blanked during the photostimulation (pink bar). **(b)** *Left:* The ΔF/F activity traces of the most excited S2 cell (of the session of panel A) are shown, averaged across stimulus conditions. This S2 cell shows a large response on hit trials (green) but no response on miss trials (red) or reward only trials (blue). The transparency of the line indicates the number of cells targeted in S1. Trials in which 150 cells were targeted were removed for display. *Right:* Equivalent plot for the most inhibited S2 cell of panel A. **(c)** The average population response to hit and miss trials across all sessions is shown (shaded areas show 95% confidence interval across trials of all sessions). Traces are averaged across cells, trials and sessions for a given trial type. Trials in which 150 cells were targeted were removed for display. The population responses of all other trial types are shown in **Extended Data Fig. 5b,c. (d)** The neural response in S1 on hit and miss trials depends on the number of cells targeted in S1. *Left:* The fraction of excited cells (**methods**) in S1 maps linearly to the number of cells targeted on both hit and miss trials. *Right:* The fraction of inhibited cells in S1 maps linearly to the number of cells targeted on both hit and miss trials. The shaded purple bar shows the 95% confidence interval across sessions of the fraction of excited or inhibited cells in S1 on reward only (R.O.) trials. The linear fit was determined using Weighted Least Squares, where the weights were the inverse variance of the trials that constituted a data point, and subsequently bound between their 25^th^ and 75^th^ percentiles to prevent extreme weight values^108^. **(e)** Equivalent panel for S1 responses. *Left:* There is no relationship between the fraction of excited cells in S2 and the number of cells targeted in S1 on hit trials or miss trials. The shaded purple bar shows the fraction of excited or inhibited cells in S2 on reward only trials. *Right:* The fraction of inhibited cells in S2 maps linearly to the number of cells photostimulated in S1 on both hit and miss trials. **(f)** The fraction of cells excited by photostimulation in S1 is highly correlated with the fraction of cells inhibited following photostimulation, both on hit trials *(left)* and miss trials *(right)*. The size of the circle indicates the number of cells photostimulated. **(g)** The fraction of cells excited by photostimulation in S2 is correlated with the fraction of cells inhibited following photostimulation, both on hit trials *(left)* and miss trials *(right)*. Significance is indicated by: *** (p < 0.001), ** (p < 0.01), * (p < 0.05) or n.s. (not significant).

We observe that hit trials elicit a significantly greater fraction of responsive cells than any other trial type in S1. (Wilcoxon signed-rank test, p < 0.05, Bonferroni corrected; **Extended Data Fig. 5a)**. In S2, the fraction of responsive cells was more similar across trial types, suggesting that stimulus information is not encoded by this relatively simple metric (**Extended Data Fig. 5a**, Wilcoxon signed-rank test, Bonferroni corrected; p < 0.05 for hit vs 2 out of 4 trial types). However, on a single cell level, we find some individual neurons in S2 that are excited or inhibited in response to hit trials, but not miss or reward only, hinting that reported stimuli are propagated to single neurons in S2 **(Fig. 2b)**.

Interestingly, however, averaging responses across cells, trials, and sessions reveals that hit trials elicit only a modest increase in population activity from baseline in both S1 and S2 (peak post-stimulus ΔF/F = 0.038 ± 0.004 and 0.022 ± 0.006, respectively) followed by a prolonged period of inhibition, while on miss trials the population average response did not deviate from baseline in S2 **(Fig. 2c)**.

The population responses of S1 and S2 suggest that excitatory stimulation is balanced by an inhibitory response **(Fig. 2a)**. The relatively weak grand average response also supports that notion **(Fig. 2c)**. Therefore, we analysed the excited and inhibited cells separately. Indeed, the inhibited responses increased with the strength of optogenetic stimulation (number of cells targeted; **Fig. 2d,e right**; correlation values reported in the figure). In S1, but not in S2, the excitatory response also increased with stimulation strength **(Fig. 2d,e left)**. This may reflect the strong contribution of the targeted cells in S1.

To quantify this effect directly, we analysed the excitation-inhibition ratio across all sessions. We find a clear correlation between the fraction of excited and inhibited cells for both hit and miss trials in S1 and S2 (**Fig. 2f,g**; correlation values reported in the figure). This indicates that the excitation-inhibition ratio was maintained regardless of photostimulation strength or whether the stimulation was perceived. However, downstream from the stimulation, in S2, the correlation between excitation and inhibition was weaker than in S1 (excitation-inhibition correlation of S1 hit versus S2 hit, p=1.2×10^−5^; S1 miss versus S2 miss, p=0.001; Two-tailed Steiger test; **Fig. 2f,g)**. Taken together, we show that as activity is propagated from its site of initiation in S1 to S2, the excitation-inhibition interplay persists but is less tight.

## S1 and S2 populations encode representations of perceived photostimulation

Individual cells displayed a diverse array of responses, and balanced population responses masked the magnitude and time course of propagated activity **(Fig. 2)**. To gain insight into what task-related information is contained in these complex neural responses, we assessed propagation of stimulus-relevant activity from S1 to S2 and its relationship to behaviour. We trained classifiers to dynamically decode the trial type of individual trials using the activity of all cells in S1 or S2 separately **(Fig. 3a)**. First, we trained classifiers that distinguish hit trials from correct rejection trials in order to detect the neural signature of both the stimulus and the activity resulting in its perception and report. Trained independently on each time frame, the classifiers performed significantly above chance for more than three seconds after stimulation, both in S1 and S2 (two-sided Wilcoxon signed-rank test, p < 0.05, Bonferroni corrected for number of tested time points; **Fig. 3b,c; Extended Data Fig. 6a,b; Extended Data Fig. 7a,b)**. This shows that neural activity underpinning perceived stimulation persisted both locally in S1 and downstream in S2 for several seconds.

**Figure 3:**
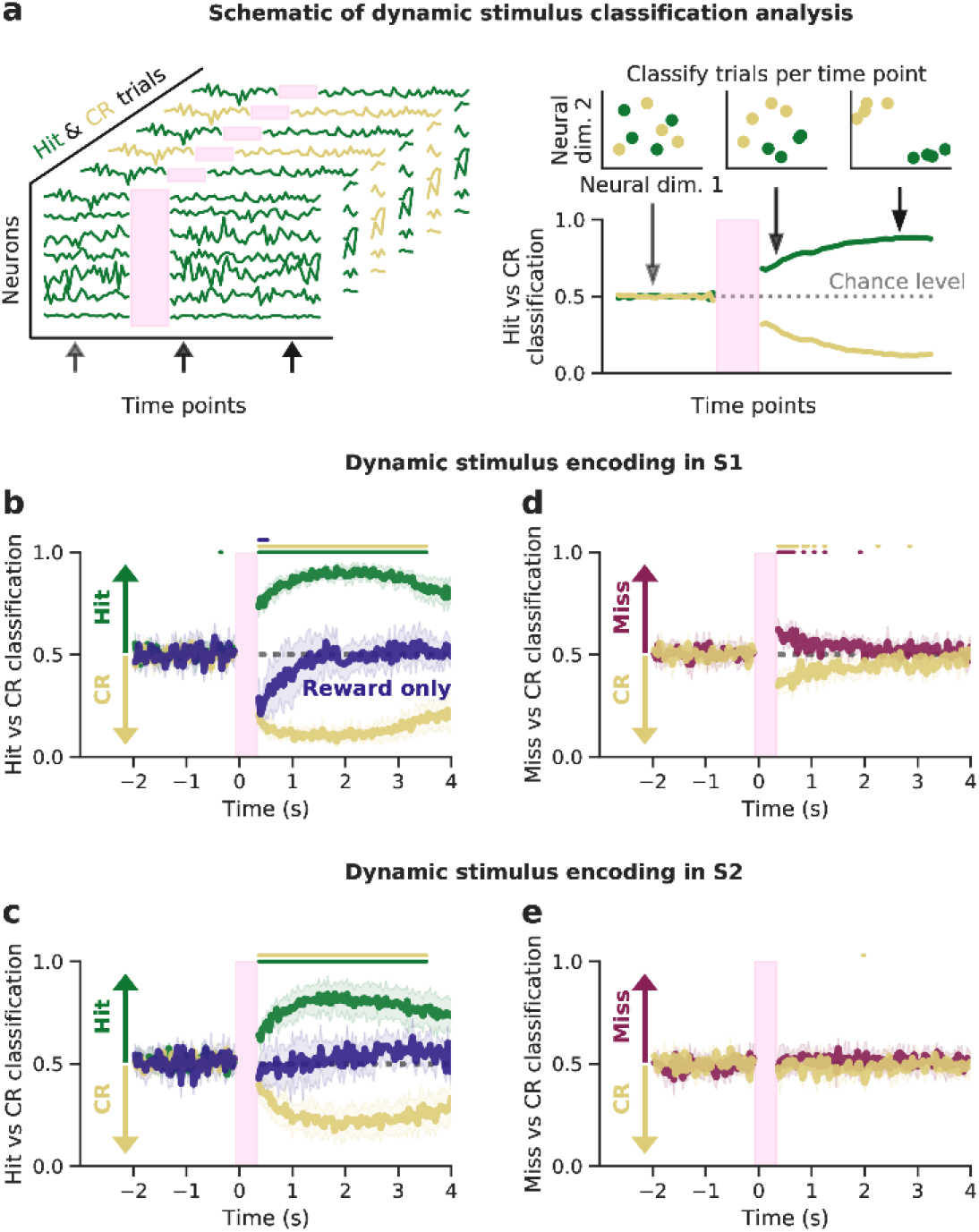
Perceived photostimuli elicit persistent activity in both S1 and S2 populations. **(a)** The strength of trial type decoding in the neural population in S1 was dynamically quantified using logistic regression classifiers. Classifiers were trained on each time frame individually, with activity of all cells in S1 or S2, and tested on held-out data. **(b)** Classifiers were trained, for each time point, on S1 activity to classify hit trials from correct rejection trials and then tested on held-out hit trials (green), correct rejection trials (yellow) and reward only trials (blue). Coloured bars above the traces show time points at which classifier performance was significantly different from chance (p < 0.05, Bonferroni corrected, see **methods**). The classifiers were able to distinguish hit trials from correct rejection trials with high accuracy for several seconds following photostimulation, implying that activity that arose from perceived stimulation persists in S1. Reward only trials were not classified as hits, showing that the classifiers were not just trained to decode the neural signature of reward on hit trials. **(c)** Classifiers were trained on S2 activity to distinguish hit trials from correct rejection trials and then tested on hit trials (green), correct rejection trials (yellow) and reward only trials (blue). As above, the classifiers were able to decode hit trials from correct rejection trials for several seconds following photostimulation, implying that activity that arose from perceived stimulation in S1 is propagated to S2 and persists for several seconds. Reward only trials were not classified as hits, indicating that the model was not just detecting the neural signature of reward on hit trials. **(d)** Classifiers were trained on S1 activity to classify miss trials from correct rejection trials and then tested on miss trials (red) and correct rejection trials (yellow). The classifiers were able to distinguish the two trial types for only for ∼1 second following photostimulation. This implies that non-perceived stimuli do not generate persistent activity. **(e)** Classifiers were trained on S2 activity to classify miss trials from correct rejection trials and then tested on miss trials (red) and correct rejection trials (yellow). The classifiers were not able to classify miss from correct rejection trials, indicating that non-perceived stimuli were not robustly propagated from S1 to S2, likely because they were also not encoded in S1 (panel **d**).

However, the classifiers could just have decoded the signatures of reward or the motor command required for licking, which are present on hit trials but not correct rejection trials, instead of decoding stimulus perception. To disambiguate this, we tested the classifiers (that had been trained to distinguish hit versus correct reject trials) on “reward only” trials (i.e. trials in which reward was delivered in the absence of photostimulation). Performance on reward only trials was either below or at chance level in S1, and remained at chance throughout the trial in S2, indicating that neural signatures of motor commands and licking were not the driving features of the classifier **(Fig. 3b,c; blue lines; Extended Data Fig. 6a,b**, p > 0.05, Bonferroni corrected).

To evaluate the neural response to reward, we trained classifiers to distinguish reward only from correct rejection trials; for these classifiers, hit trials were evaluated at equal accuracy to reward only trials (**Extended Data Fig. 6e-h**, dark blue and green traces). This indicates that the neural response related to reward is comparable for both trial types, and that neural signatures of perceived stimulation are present on hit trials, both in S1 and S2.

Next we asked whether neural activity from unperceived photostimulation was reliably propagated from S1 to S2 by training models to classify miss and correct rejection trials (**Fig. 3d,e; Extended Data Fig. 5c,d; Extended Data Fig. 7c,d)**. We found that in S1, these classifiers decoded miss trials slightly above chance immediately following stimulation (p < 0.05, Bonferroni corrected), however performance rapidly decayed back to chance level after 1 second. Conversely in S2, the classifiers were not able to decode miss trials, indicating that stimulus information that is not perceived does not propagate to S2.

Taken together, these results show that only perceived stimulus representations are reliably propagated out of the local brain region in which they originate to a downstream brain area. Further, the representations of stimuli that are propagated and drive perception persist both locally and downstream for several seconds following the injection of activity.

## Pre-stimulus S1 activity predicts trial outcome

While the previous analysis clearly showed that stimulation of S1 can be reliably propagated to S2 and reported behaviourally, it is unclear what intrinsic network conditions facilitate its detection. To identify the conditions that facilitate both the propagation to S2 and the behavioural report, we analysed population activity in S1 immediately *prior* to stimulation **(Fig. 4a)** and asked whether the pre-stimulus population activity could predict whether the stimulus would be perceived (i.e. whether the upcoming trial type would be a hit or a miss). To characterise population activity, we analysed the distribution of instantaneous activity across neurons and computed the population mean and population variance as scalar measures. The mean population activity was not predictive of whether a trial will be a hit or miss (p=0.28, Wilcoxon signed-rank test, **Fig. 4b left**). In contrast, the population variance across S1 neurons strongly predicted whether the upcoming stimulus would be a hit or miss trial. The population variance in S1 was larger prior to miss trials than prior to hit trials (p=0.001, Wilcoxon signed-rank test, **Fig. 4b right**). We further observed that population variance was correlated with task variables such as trial number and reward history **(Extended Data Fig. 8)**, and that population variance in S1 was correlated with that of S2 **(Extended Data Fig. 8e)**. This suggests that population variance in both, S1 and S2, may be a global mechanism underpinning how arousal and/or motivational state are instantiated in neural circuitry. The relationship between population variance and behavioural performance was also evident on a trial-by-trial level, whereby increased population variance pre-stimulus in S1 is negatively correlated with the probability that the upcoming stimulus elicits a hit (**Fig. 4c;** one-sided regression Bonferroni corrected, test p < 0.001 on 10/11 sessions). Further, even though the population variance influenced the probability of the stimulus being perceived (and thus propagated downstream) (**Fig. 4c**), the subsequent strength of stimulus encoding was not influenced by population variance (**Extended Data Fig. 9**). This implies that the neural representation of perceived stimuli was independent of the preceding population variance, and therefore generalised.

**Figure 4:**
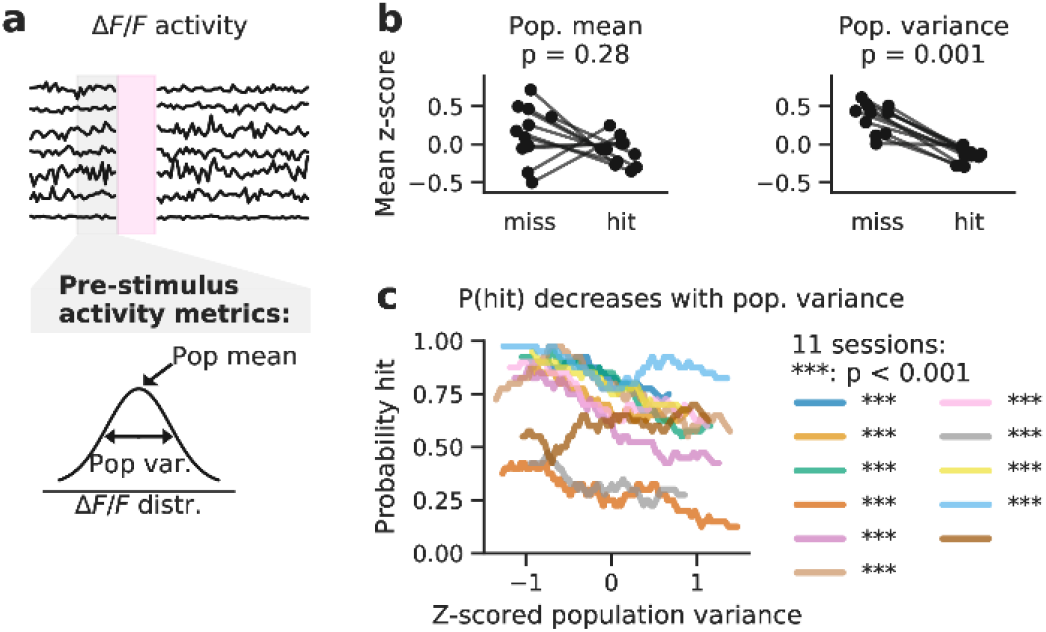
Pre-stimulus population in S1 predicts the upcoming trial outcome. **(a)** Illustration of neural activity throughout a trial. Only the activity in the 0.5 seconds before the stimulus on a given trial is included in subsequent panels. First, we considered 2 metrics of pre-stimulus neural activity; the population mean and population variance. **(b)** Comparison of population metrics of pre-stimulus S1 activity prior to hit trials and prior to miss trials. *Left*: No evidence that mean population activity pre-stimulus predicts the upcoming trial outcome. *Right*: Population variance is significantly higher before miss trials than before hit trials. Pre-stimulus population metrics in S1 were computed trial-wise and z-scored across trials within a session before being split into hit and miss trials and averaged across a session. P values tested for a difference in session-wise population metrics between hit and miss trials (Wilcoxon signed-rank test). **(c)** The probability of detecting the photostimulation decreased linearly with increasing pre-stimulus population variance in S1 for 10/11 sessions. Trials within a session were binned by their z-scored population variance and this was correlated to the probability of a hit trial within that bin.

## Strong pre-stimulus recurrent interactions prior to miss trials

Measuring population mean and variance immediately pre-stimulus is the most direct way of quantifying how instantaneous activity impacts perception **(Fig. 4)**. However, more time points are needed to quantify the covariance structure, which could link these basic measures of population activity to mechanistic theories of circuit function. Therefore we concatenated the activity that preceded all miss trials and hit trials separately, and assessed their differences **(Methods, Extended Data Fig. 10a-c)**. We explored two complementary analytical approaches to characterise the role of population activity in driving behaviour.

First, the strength and structure of inter-neuronal correlations^47^ can affect sensory-guided behaviour^48,49^. Hence we assessed whether the low-dimensional correlation structure of pre-stimulus population activity, measured as the variance explained by the first 5 latent factors, could predict the perceptual variability we observed, but found no evidence of this in our experiments (**Fig. 5b-c**, p=0.168, two-sided Wilcoxon signed-rank test). We also tested whether the average “non-shared” pairwise correlation of pre-stimulus activity was predictive, but again found no evidence for this hypothesis (**Fig. 5d-e**, p=0.365, two-sided Wilcoxon signed-rank test).

**Figure 5:**
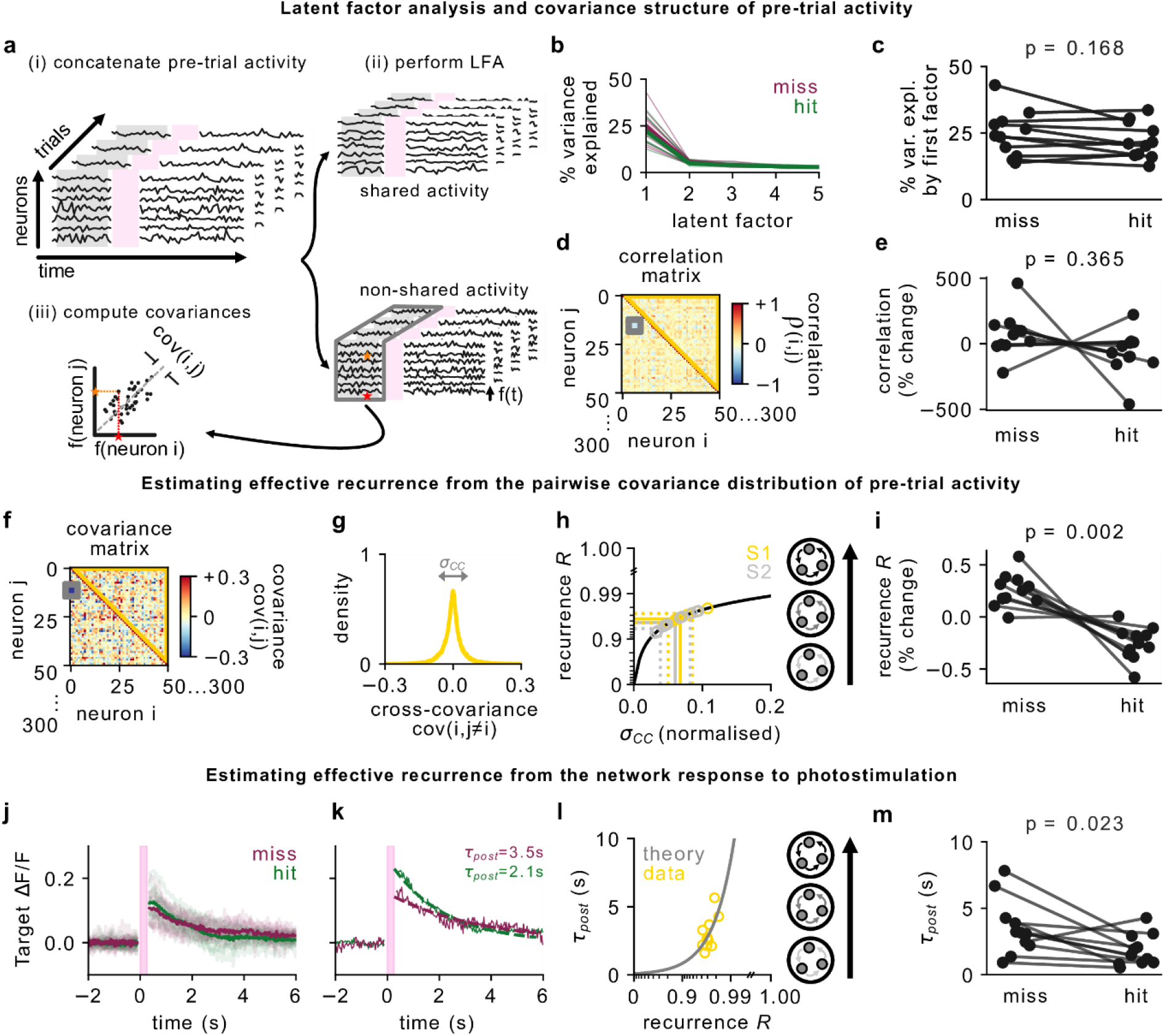
Miss trials are preceded by higher effective recurrence. **(a)** To quantify effective recurrence *R* in a network, we first apply latent factor analysis to identify activity that is putatively driven by shared external input. We then subtract out this “shared” activity and focus on the remaining “non-shared” activity. **(b)** Variance explained by each of the first 5 latent factors (miss: red, hit: green). **(c)** The variance explained by the first factor (also called “across-neuron population-wise correlation”^48^) is not significantly different between hit and miss trials (two-sided Wilcoxon signed-rank test, p=0.168). **(d)** For 6.5 seconds of non-shared pre-trial activity, the cross-neuron correlation matrix is calculated (orange and red stars show the process for a pair of example neurons, grey box highlights that of a single pair, yellow triangle marks the off-diagonal entries that are analysed further). **(e)** The mean off-diagonal cross-correlation is not significantly different prior to hit or miss trials (p=0.37, two-sided Wilcoxon signed-rank test). **(f)** Cross-covariance matrix of the non-shared activity (yellow triangle marks the off-diagonal entries used to compute recurrence). **(g)** Histogram of the cross-covariance matrix, grey arrows indicate the variance of the distribution *σ*_*CC*_. **(h)** The relationship between *σ*_*CC*_ and the effective recurrence *R* of the local network is known from theoretical derivations (see Methods, plotted in black for a network of 50,000 neurons). Data for individual sessions is shown as gold and silver circles (S1 and S2, respectively), straight lines the average across sessions, and dotted lines show the spread (mean ± std). **(i)** The effective recurrence *R* prior to stimulation is significantly different on hit and miss trials, suggesting that lower recurrence facilitates stimulus detection (p=0.002, two-sided Wilcoxon signed-rank test). **(j)** The average photostimulation response of the targeted cells on either hit or miss trials (excluding trials where 150 targets were stimulated, averaged across sessions in bold and averaged per session in shade). **(k)** The “network response timescale” *τ*_*post*_ was determined by fitting an exponential decay function per session. **(l)** The inferred *τ*_*post*_ values (yellow circles) were better explained by the linear network theory (grey line, *r*^2^ = 0.44) than a simple linear regression (not shown, *r*^2^ = 0.38). **(m)** The inferred network response timescale *τ*_*post*_ is significantly different on hit and miss trials (p=0.023, two-sided Wilcoxon signed-rank test).

Second, neural circuits are characterised by strong recurrence^50–52^, which is known to give rise to trial-to-trial variability^53^; moreover, the strength of recurrence can determine local amplification and encoding of external stimuli^54–57^. We inferred the strength of recurrence in the local network (*R*) from the non-shared covariance distribution as previously described^57^ **(Fig. 5f-h, Extended Data Fig. 10)**, and found that neural dynamics in both S1 and S2 are strongly recurrent (*R* = 0.964 ± 0.008 in S1, *R* = 0.957 ± 0.016 in S2, **Fig. 5h**), similar to other cortical areas in rodents^51,57,58^. Importantly, we also observed that lower recurrence *R* prior to stimulation predicts higher detection probability (p=0.002 in S1 **Fig. 5i, Extended Data Fig. 11o, Extended Data Fig. 12i,j** and p=0.050 in S2, **Extended Data Fig. 11r**, two-sided Wilcoxon signed-rank tests). We next compared this network-wide, covariance-based recurrence metric to the network response timescale *τ*_*post*_, a measure of coupling strength inferred directly from the activity of the photostimulated targets^59^, and found the same trend as for the recurrence *R* (**Fig. 5j-m**, p=0.023, two-sided Wilcoxon signed-rank test). These data suggest that the previously described trial-by-trial differences in population variance might result from changes in the effective recurrence strength prior to stimulation, influencing the network’s sensitivity to direct stimulation.

## Stimulus strength and neural population variance determine probability of perception

Our previous analyses show that we have identified two separate conditions that predict the probability a stimulus will be perceived: the strength of the stimulus (number of cells targeted, **Fig. 1f**), and the variance of the population activity (**Fig. 4b,c**). Next we asked if these two predictors work in tandem, i.e. whether stimulus perception depends on both the strength of the stimulus and the ongoing state of the population.

We find that the probability that a stimulus elicits a hit can be expressed on a two-dimensional axis **(Fig. 6a)**, where hit probability is highest when population variance is minimised and the number of cells targeted is maximised. More targeted cells were required to reliably drive hits, as pre-stimulus population variance increased. This phenomenon can be conceptualised by a signal-to-noise ratio (SNR) framework, in which activity injected into cells through photostimulation forms the signal and the magnitude of background noise is measured by the population variance. The higher the SNR, the more likely the animal is to detect the signal above ongoing noise and respond, as quantified by collapsing the population variance and the number of cells targeted onto a single SNR axis (**Fig. 6b**, both population variance and the number of targeted cells contributed significantly to predicting trial outcome, both p < 10^−13^, logistic regression two-sided t-test, see Methods). Further, as pupil size changes have been associated with spontaneous activity fluctuations^13^, we measured the change in pupil size of 3 animals during the all-optical experiments, but found no evidence that pre-stimulus pupil size influenced trial outcome (**Extended Data Fig. 14**). In summary, our results show that the SNR of a stimulus determines how likely it is to be perceived and propagated downstream to drive behaviour **(Fig. 6c)**.

**Figure 6:**
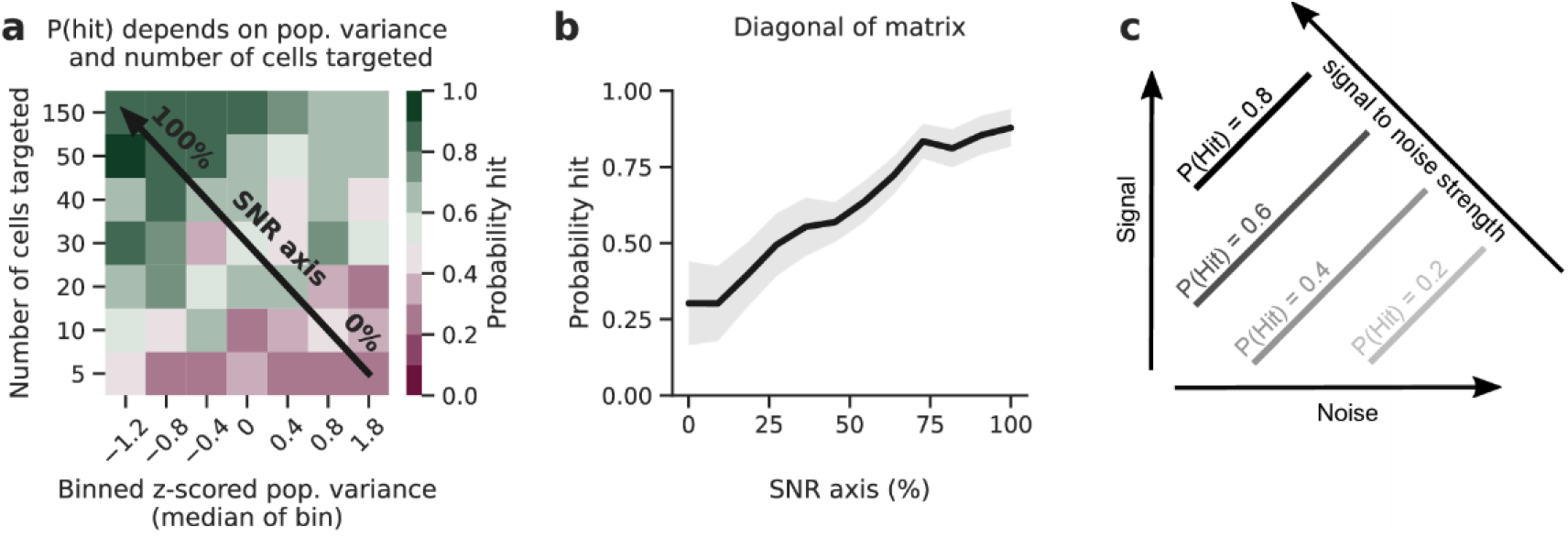
Stimulus strength and neural population variance underpin perception. **(a)** The interaction between pre-stimulus population variance in S1 and the number of cells targeted by photostimulation defines the probability of a hit trial. Trials were binned by their z-scored population variance and by the number of cells targeted; the probability of a hit within each bin is plotted on a 2-dimensional axis, pooled across all sessions Increased the number of cells targeted (i.e. signal strength) and decreased the pre-stimulus variance (i.e. ‘noise’) generally yielded a greater probability of a hit trial. This is referred to as the Signal to Noise Ratio (SNR), as indicated by the diagonal black arrow. **(b)** Maximising the SNR of the stimulus resulted in the maximal probability of a hit trial. Data as in **a**, but projected onto the SNR axis - as indicated in panel **a** - by averaging across all bins that project orthogonally onto each point on this axis. **(c)** Schematic outlining the intuition for the SNR axis. Increasing the number of cells targeted on a given trial maximises the signal of that stimulus. Noise is proportional to the population variance as there is more excitation and suppression from baseline in a population with high variance. The probability of hit is maximal when SNR is maximal, as the stimulus is more likely to be detected above ongoing activity.

## Discussion

Critical to the brain’s ability to process stimuli is that activity is robustly propagated between functionally and anatomically distinct regions^60–62^. While this question is often addressed in observational meso-scale studies of particularly the primate brain^63,64^, less is known about how the dynamics of neural activity are structured at the single-cell level to propagate activity between brain regions (but see^19,20,65^). By using an all-optical interrogation technique across multiple brain areas for the first time, we demonstrate that behaviourally salient photostimulation of S1 as a causal intervention elicits robust activation of S2. As expected, stronger stimulation elicits a stronger response in S2. Interestingly, we also find that pre-stimulus activity in S1 influences the detectability of the upcoming stimulus. This is consistent with a signal-to-noise (SNR) framework in which the signal strength is the number of photostimulated cells and the noise is the pre-stimulus neural variability in S1. Thus the SNR of an input relative to the network state is critical both for its behavioural salience and the likelihood that its representation is propagated beyond the brain region of origin.

One feature of cortical dynamics that may allow activity to reliably propagate is inhibition stabilisation in which inhibition tracks excitation to prevent unstable dynamics^66–69^. Indeed, we observed that excitation and inhibition covaried with stimulus strength in S1 and S2, and that both regions demonstrated correlated excitation and inhibition. Taken together, our results show that the population response of somatosensory cortex to excitatory photostimulation of small groups of neurons was stabilised by inhibition both in the excited region and in a downstream region. We observed that both the correlation between excitation and inhibition, as well as the fraction of inhibited cells, were lower in S2 than S1. This could explain why stimulus decodability was also slightly better in S1, as theoretical results suggest that strong inhibition enhances stimulus representations ^70^.

Further, we found that the population representation of the stimulus was generalised, as the neural response to perceived stimulation was comparable across trials in both regions. We demonstrated this by showing that classifiers robustly decoded hit trials using cross-validation *across* trials, even though we varied the identity of targeted neurons on *each* trial. This variation of the stimulus requires sufficient consistency of the representation of the stimulus, implying that perceived neural activity remains generalised throughout its journey through the cortex. Conversely, on miss trials, the presence of the stimulus was only decodable for a brief period in S1 immediately following stimulation, and not at all in S2, showing that non-perceived stimulus information was not generalised across both brain regions. Stimulus generalisation is a well-characterised phenomenon in psychology^71,72^, biological circuits^73,74^ and in artificial neural networks^75–77^, and endows an agent with the ability to effectively interpret novel stimuli based on prior experience. This process has been shown to be enhanced if the stimulus is coupled to reward^74^, matching our results.

The SNR of a sensory neuron^78^ or population of neurons^79^ is often used to quantify the fidelity of the representation of a stimulus, whereby a higher SNR means that the sensory stimulus is more robustly represented in neural activity. Indeed, one of the functions of the highly recurrent circuitry in sensory cortex is thought to be the amplification of relevant activity arising from feedforward inputs, in order to enhance SNR^19,80,81^. Despite the well-characterised importance of high SNR in the local representation of sensory stimuli, it is unclear how the SNR of a stimulus relates to its likelihood to propagate downstream. Here we generated a signal with direct cortical activation and quantified noise using the population variance metric. We found that the probability a photostimulus was propagated and perceived was determined by the SNR of the stimulus.

Other metrics of pre-stimulus neural variability, such as single-cell temporal variance^13,82^, synchrony^16,83^ and neural oscillation power^19,84,85^ have all previously been linked to task performance. Although differences in recording techniques preclude direct comparisons, in particular, lower variance of single-cell membrane potentials were also found to enhance stimulus detection performance^82,86^, possibly providing a link between single-cell and population variability. Population variance thus captures the population-wide neural variability, which can arise from recurrent neural activity^57^.

We quantified the recurrence *R* in order to test theories^80,87,88^ about its impact on signal propagation. We found that the S1 network operates in a strongly recurrent dynamical regime, similar to other cortical areas in rodents^51,57,58^, monkeys^52,89,90^ and humans^91^, in which a small change in *R* can have a large effect^87,92^. Indeed, we also found that slight variations around this operating point prior to stimulation are predictive of detection performance. Specifically, we found that higher recurrency was associated with a decrease in stimulus detection. This is consistent with the SNR framework outlined above whereby greater noise (variability) results in diminished perception, and with previous results that show that recurrence can be tuned to meet task requirements^52,89–91^. Therefore, the causal nature of our experiment allowed us to extend previous work by directly injecting a controlled stimulus while observing the neural activity, thereby linking changes in recurrence to perception.

We claim that the photostimulus is causal in driving both behaviour and observed neural responses. Photostimulation was designed such that it causally drove behaviour as the task required the mouse to lick following photostimulation to receive a reward. Our data confirm that indeed, the probability of licking immediately following stimulation was far greater than baseline. The causal paths that lead to the observed neural activity, in particular the transmission to S2, is less trivial because in practice, causality is challenging to define^93^ or to quantify in the context of neural activity recordings^94^. One crucial criterion for causality generally but particularly important in neuroscience is that an effect should consistently follow a randomised intervention^97^. In our experiments, neural responses to perceived photostimulation in S2 were sufficiently consistent despite the random timing of their delivery such that we could classify them as specific to the Hit condition. This resistance to randomisation of the intervention critically underpins our argument for causality. Furthermore, our observed effects also satisfy some of the properties of observational causality (a weaker form of causality), namely strength (effect size), consistency (effect reliably follows stimulus), temporality (effect follows stimulus closely in time), dose-response relationship (greater stimulus leads to greater effect), and plausibility (reasonable mechanistic explanation)^93–97^. The strength, consistency, temporality, and dose-response relationship of neural activity in S2 following S1 photostimulation are shown in **Fig. 2, 3, 6** and **Extended Data Fig. 13**. The information transmission from stimulation to S2 neurons could be caused via many different paths. We do not have the experimental capacity to disentangle all potential paths driving S2 activity. These include, but are not limited to: direct, monosynaptic transmission from S1 to S2^38,41,98^, indirect paths via other brain regions^99^, brain state changes due to reward consumption^83^, or reafference from the whiskers^100^ or muscles^95,96^. Thus, while we can make statements about a causal relation, we cannot make any claim about the specific paths or their relative contributions.

Theoretical work has also highlighted the potential role of feedback connections in information processing^101–104^, and it is known that S2 also projects back to S1 directly and indirectly^38,41,98,105,106^. An interventional approach e.g.^23^ could further interrogate the role feedback in S1-S2 circuit, to elucidate whether perception arises through a bi-directional flow of information.

In summary, our study examines for the first time how injecting activity into small groups of neurons in an upstream brain region impacts both behaviour and propagation of activity to a downstream region. We find that sufficiently strong stimulation of somatosensory cortex, relative to pre-stimulus neural variability, elicits generalised responses that are inhibition-stabilised and propagate downstream to drive behaviour. Therefore, building on rigorous analysis of our data, we outline a signal-to-noise framework linking causal activation to perception.

## Methods

All experimental procedures involving animals were conducted in accordance with the UK animals in Scientific Procedures Act (1986).

Male and female C57/BL6 and Tg(tetO-GCaMP6s)2Niell mice were used for all experiments. Mice were between 4-12 weeks of age when surgery was performed.

### Data and software availability

The functional two-photon calcium imaging recordings that are presented in this manuscript will be made publicly available in a GIN repository upon publication.

Behavioural training used pyControl hardware and software as previously reported^109^. Photostimulation was controlled by custom written code in Python and C, available from the authors upon request. Online imaging analysis used STAMovieMaker (https://github.com/llerussell/STAMovieMaker). Offline pre-processing imaging analysis was performed using Suite2p^110^.

All subsequent data analysis and visualisation was performed in Python 3.7 using custom written code, which will be made publicly available on Github upon publication.

### Surgical Procedures

Animals were anaesthetised with isoflurane (5% for induction, 1.5% for maintenance) during all surgical procedures. A perioperative injection of 0.1 mg/kg buprenorphine (Vetergesic), 5 mg/kg meloxicam (Metacam) was administered. Mice were prepared for chronic imaging experiments through a single surgery. 2 mg/kg Bupivacaine (Marcaine) was applied to the scalp before it was sterilised with chlorhexidine gluconate and isopropyl alcohol (ChloraPrep) before being removed bilaterally. The skull was cleaned with a bone scraper (Fine Science Tools) to remove the periosteum. An aluminium head-plate with a 7 mm imaging well was bonded to the skull using dental cement (Super-Bond C&B, Sun-Medical). A 3 mm circular craniotomy was drilled over the right somatosensory cortex, targeting the S1/S2 border (-1.9 mm posterior, +3.8 mm lateral), using a dental drill (NSK UK Ltd.). The skull within the craniotomy was soaked in saline before removal. Any blood was flushed with saline for >5 minutes, before a durotomy was performed. A single 1 μl viral injection was performed using a calibrated injection pipette bevelled to a sharp point. Injections were performed at a rate of 100 nl / minute at 300 μm below the pial surface and were controlled using a hydraulic micromanipulator (Narishige).

Pipettes were front loaded with either: 1:10 GCaMP6s (AAV1-Syn.GCaMP6s.WPRE.SV40) diluted in C1V1-Kv2.1 (AAV9-CamKIIa-C1V1(t/t)-mScarlet-KV2.1) if injecting into C57/BL6 mice or C1V1-Kv2.1 alone if injecting into transgenic mice. After injection, a double tiered cranial window composed of a 4 mm circular coverslip glued to a 3 mm circular coverslip was pressed into the craniotomy and sealed with cyanoacrylate (VetBond) and dental cement. Mice were recovered in a heated recovery chamber and kept under observation until behaving normally. Mice were subsequently monitored and their weight recorded for 7 days following surgery. Mice were allowed to recover for at least 21 days with *ad libitum* access to food and water before further procedures. This also allowed viral expression to ramp up before behavioural training was commenced.

### Two-photon imaging

Two-photon imaging was performed using a resonant scanning microscope (2PPlus, Bruker Corporation) which raster scanned a femtosecond pulsed, dispersion-corrected laser beam (Vision-S, Coherent) across the sample at 30 Hz. A 16x/0.8-NA water immersion objective lens (Nikon) was used. GCaMP and mScarlet were imaged using a 920 nm and 765 nm beam respectively. Power on sample was controlled using a Pockels cell (Conoptics) and was kept at 50 mW for all experiments. A rectangular field of view (1024 × 514 pixels, 1397.4 × 701.4 μm), was used to image across two brain regions at 30 Hz. Imaging was controlled through PrairieView (Bruker Corporation).

### Widefield imaging

Whisker stimulation during widefield calcium imaging was used to: target virus/tracer injections, determine the viability of a preparation for continued experimentation and find a suitable field of view for two-photon imaging encompassing S1 and S2 responses to a single whisker deflection. Using a camera, an LED and a dichroic/filter set, the whole cranial window was imaged using epifluorescence to measure calcium responses to whisker deflections (**Extended Data Fig. 1)**. Each one of four whiskers (B1-B3 and C2; one at a time) was threaded into a capillary and deflected for 10 trials of 1 second each with 10 second inter-trial intervals. Each stimulation was 10 Hz and of ∼300 um in amplitude at ∼500 um from the base of the follicle. Responses to whisker stimulation were assessed using ΔF/F stimulus-triggered averages based on a baseline of 1 second pre-stimulus. Two areas of response moved stereotypically anterior-posterior and medial-lateral according to the row and column of the whiskers stimulated, confirming whisker responses were being assessed (**Extended Data Fig. 1b, c**).

### Pupil Imaging Analysis

Facial videography was performed from the left side of the animal, on sessions 7, 8 and 9 (corresponding to Extended Data Fig. 4). Eyes were illuminated using an infrared light source (Amazon, 48 LED illuminator light), and videos were recorded at 30 Hz (each frame triggered by the 2p microscope control software to ensure synchronicity) and 560×500 pixel (8 bit resolution) with the camera (Point Grey CM3-U3-13Y3M-CS) placed close to the mouse.

For analysis, videos were cropped around the eye (mean values: 164 pixels in x and 140 pixels in y, eye diameter is 122 pixels). For each video, the pixel intensity histogram of each frame was normalised to match contrast levels. Next, to segment the pupil, frames were binarised so all pixels with an intensity below a threshold value (10/255 for 2 sessions, 12/255 for 1 session) were 1 and all other pixels were 0. This sparsely labels the pupil while keeping false positives outside of it to a minimum. The best-fitting ellipse covering all pixels of value 1 around a fixed centroid (manually annotated once per video) was then selected via a Nelder-Mead simplex algorithm^111^, implemented using the *scipy*.*optimize*.*minimize* function^112^ with L0 penalty. For each frame, the major axis of the ellipse was used as an estimate of pupil radius, and a median filter with a window of 6 frames (implemented in the *scipy*.*ndimage*.*median_filter* function) was applied to eliminate outliers. Finally, for the analysis of pupil size on hit and miss trials (Extended Data Fig. 14), pupil radius was normalised, per session, by subtracting the average pupil size radius in the 2 seconds pre-stimulus across all hit and miss trials.

### Two-photon optogenetic stimulation

Two-photon optogenetic stimulation was conducted using a pulsed fixed-wavelength fibre laser at either 1030 nm (Satsuma HP2, Amplitude Systèmes) or 1035 nm (Monaco, Coherent) at a repetition rate of 2 MHz. Multiple individual neurons were targeted for stimulation simultaneously by splitting the laser beam using a reflective spatial light modulator (SLM) (7.68 × 7.68 mm active area, 512 × 512 pixels, Boulder Nonlinear Systems). The active area of the SLM was overfilled and the polarisation optimised for maximal first order diffraction efficiency using a half-wave plate. The zero order diffraction beam was blocked using a well bored into an optical flat using a dental drill (NSK UK Ltd).

Phase masks were loaded onto the SLM using the Blink SDK (Meadowlark Optics). Phase masks were computed by applying the Gerchberg-Saxton algorithm^113^ to the xy coordinates of the target cell bodies. A weighted form of this algorithm was used to ensure uniform power distribution across all cells as the first order diffraction efficiency of the SLM is reduced with increasing distance from the zero order location. An image of the SLM was relayed onto a pair of galvanometer mirrors (Bruker Corporation) integrated into the two-photon imaging system. The galvanometer mirrors were programmed to generate spirals across cell somata by moving each beamlet simultaneously. Neurons were stimulated using 10 × 25 ms spirals of 15 μm diameter and 6 mW power / spiral.

The affine transformation required to map coordinates from SLM space to imaging space was computed through a custom-modified version of open-source software written in MATLAB (github.com/llerussell/Naparm) using the two-photon image created by burning arbitrary patterns into a fluorescent plastic slide. Phase masks and galvanometer voltages required to perform photostimulation were generated using Naparm^114^. Voltages were applied to the galvanometers using PrairieView (Bruker Corporation). Custom written Python code was used to: select the cells for photostimulation, interface with Naparm to generate the files required to perform stimulation, interface with PrairieView to load voltage onto galvanometers and trigger photostimulation based on behavioural cues. A USB data acquisition card (National Instruments) running PACKIO^115^, was used as a master synchroniser to record the frame clock of the imaging, onset of photostimulation and a pulse to synchronise imaging and photostimulation with behaviour.

### Behavioural training

Mice were water restricted and given access to ∼1 ml of water per day. Their weights were recorded and *ad libitum* access to water or wet mash was provided if the animal’s weight dropped below 80% of the pre-restriction weight. For training, mice were head-fixed using their head-plate with their body supported in a 3d printed polylactic acid (PLA) tube. Mice became acclimatised to head-fixation and relaxed in the tube after the first 1-2 sessions.

All behavioural training was controlled using the pyControl hardware and software^116^ based around the micropython microcontroller. The pyControl framework acted as the master clock for behaviour by writing the timing of behavioural input and output events to disk and triggering trials and stimuli based on behavioural events.

Mice reported photostimulation by licking a metallic lick spout placed ∼5 mm from the tongue using a micromanipulator arm (Noga Engineering). The spout was electrically connected to the pyControl lickometer circuit (Open-ephys), which both recorded licking events and drove a solenoid valve (Lee Products) to deliver a ∼2 μl water reward.

The general structure of the task and individual trials was consistent at all stages of training. Each trial was separated by a fixed five second inter-trial-interval followed by a 4-6 second lick withhold period, where the length of the lick-withhold period was drawn randomly from a uniform distribution spanning these times. This prevented mice learning temporal structure in the task and eliminated the utility of a strategy based around random high frequency licking. If the mouse licked during the lick-withhold period, the trial was restarted and a new withhold length was drawn from the uniform distribution.

On trials where photostimulation was delivered, mice were rewarded if they licked to report perception of the stimulus. When no photostimulation was delivered, no punishment was administered for licking. No cues were made available that could signal the start of a trial.

The ‘response period’ during which the mouse’s licking response was recorded commenced immediately following the end of the lick-withhold period. This coincided with the onset of photostimulation in the case of go trials. The response period lasted for 1 second, and licks during this period alone were used to define the outcome of the trial. If the animal licked during the response period, this was scored as a ‘hit’; failure to lick on a go trial was scored as a ‘miss’. On catch trials, if the animal licked in the response period, the trial was scored as a ‘false positive’; trials where the animal did not lick in this period were scored as a ‘correct rejection’. A reward was delivered immediately after a correct lick on hit trials. This behaviour can thus be considered a detection task where catch trials are used to report the animal’s baseline licking probability. Trial type was selected pseudorandomly ensuring no more than 3 consecutive trials of the same type. Mice were trained until they ignored 10 consecutive rewards or until 90 minutes had elapsed.

Behavioural performance was quantified using the d’ metric^117^ which quantifies the difference in response probability between hit and catch trials while controlling for baseline response rate. This allows for comparison of performance of mice with conservative licking strategies, who are less likely to lick on both go and catch trials, with mice that are more likely to lick on both trial types.

d′ is defined as:

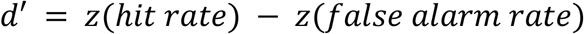

where: *z* is the Z-transform function

Sessions were discarded from all analyses if behavioural performance was poor, where d’ for trials on which 150 cells were targeted was less than 0.95 and/or d’ for trials on which 40 and 50 cells were targeted was less than 0.5.

### One-photon behavioural training

Naive mice initially learned the association between photostimulation and reward through ‘one-photon’ widefield stimulation with a 595 nm LED (Cree). One-photon behavioural training was carried out in closed, light-proofed wooden boxes with the LED placed directly onto the cranial window. Current was supplied to the LED using pyControl and a custom designed analog driver board, allowing the power produced by the LED to be controlled programmatically. LED power was calibrated using a power meter (ThorLabs PM100D). A single trial’s photostimulation consisted of 5 × 20 ms pulses over a period of 200 ms. LED stimulation was delivered on go trials only.

Initially, mice learned the association between optogenetic stimulation and reward through conditioning in which the stimulus was paired with reward regardless of the animal’s response. Rewards that were delivered on miss trials are termed ‘auto-rewards’ and were delivered at the end of the response period. Mice rapidly learned to associate the optogenetic stimulus with reward and began licking before the auto-reward within 1-2 sessions. Mice were transitioned out of the auto-reward phase and into active training when they scored three consecutive hit trials. During active training (including subsequent two-photon behavioural training, see below), mice were not auto-rewarded aside from on a single trial if they registered 3 consecutive misses; this functioned to motivate poorly performing mice during a session. These auto-rewarded trials were excluded from all further analyses. On rare occasions (33 out of 1043 total hit trials), lick responses just before the end of the response window (928 - 985 ms interquartile range from photostimulus onset) were registered just outside the response window and therefore not rewarded. These were also excluded from all further analyses.

During the initial training phase and the first stage requiring active responses, LED power of 10 mW was used, with mice transitioned to progressively weaker LED powers once their performance was sufficiently high. Online performance was computed using running d’, computed across a window of 10 trials. Once animals had reached a running d’ of 2, the LED power was reduced in a stepwise fashion, and the running window was reset, until the lowest power (0.1 mw) was reached. Once animals had exceeded criterion at this power, they were considered to have completed the one-photon training paradigm.

### Two-photon behavioural training

After learning the one-photon stimulation task, mice were transitioned to the two-photon version of the task, whereby mice responded to two-photon photostimulation targeted to S1 only. Initially mice were trained on a task in which ∼150 S1 neurons were photostimulated on every go trial in three groups of 50, with an inter-group-interval of 5 ms (each group stimulated with 10 × 25 ms spirals; entire 150 cell photostimulation takes 760 ms). Once mice registered a d’ > 1.5 across an entire session, they were transitioned to the main version of the task. This task consisted of three trial types selected pseudorandomly, with equal probability and with no more than three consecutive trials of the same type. On 1/3 of trials, 150 cells were targeted in groups of three, on 1/3 of trials, cells were targeted in a single group, with the number of targets drawn randomly from the set {5,10,20,30,40,50} with equal probability and with replacement, the final 1/3 of trials were catch trials in which no photostimulation was performed. The 5-50 target photostimulation took 250 ms (10 × 25 ms spirals).

Before each session, photoresponsive cells were identified by performing two-photon photostimulation spanning opsin-expressing areas of S1 (e.g. **Fig. 1c**). This generated the coordinates of ∼150 S1 neurons known to be responsive to stimulation. The subset of neurons to be targeted was selected randomly before each trial; cells in each simultaneously targeted subset were no more than 350 μm apart.

Prior to active behaviour, 10 minutes of spontaneous imaging was performed without any photostimulation being delivered. During this time period 10 rewards were delivered with an inter-reward-interval of 10 seconds; this allowed us to assess the ‘reward only’ neural response in somatosensory cortex. Active behaviour followed spontaneous imaging, during which the mouse was rewarded only if it responded to a go trial. Neural activity was imaged throughout active behaviour but was stopped every ∼15 minutes to ensure the objective lens was completely immersed in water and to monitor animal welfare. The field of view was manually corrected for drift throughout the session by moving the objective to realign the field of view to a marker cell.

White noise was played to the animal throughout the session to mask auditory cues signifying the onset of stimulation and galvanometer mirrors were moved in an identical fashion on both go and catch trials. This ensured that the auditory cues generated were matched on both go and catch trials, and ensured that mice were responding to optical activation of S1 alone. Behavioural events were recorded through pyControl and photostimulation was controlled by custom written routines in Python and C.

### Imaging data analysis

Online analysis carried out during an experiment was conducted using STAMovieMaker (https://github.com/llerussell/STAMovieMaker). These stimulus triggered average (STA) images displayed visually the change in activity of each pixel in the field of view post-stimulus relative to pre-stimulus, thus the pixel intensity of the image was proportional to the increase in activity driven by photostimulation. STA images were trial averaged in a group-wise fashion and coloured, such that pixels of a given colour represent the average activity driven by repeated photostimulation of a given group of neurons. These images were used to manually define the coordinates of cells in S1 responsive to optical stimulation.

All further analysis was conducted offline. Calcium imaging movies were processed using Suite2p^110^ and regions of interest (ROIs) corresponding to putative cell somata were manually selected. Suite2p also extracts a signal arising from the neuropil surrounding a cell body. To remove contamination of the signal arising from individual soma by the surrounding neuropil, we subtracted the neuropil signal from each cell body at each timepoint (t) according to the equation:

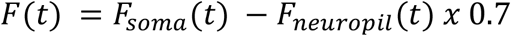

where:

*F* = neuropil subtracted fluorescence

*F*_*soma*_ = fluorescence from the cell’s soma

*F*_*neuropil*_ = fluorescence from the cell’s surrounding neuropil

0.7 = neuropil coefficient^45^

To ensure that cells with a bright baseline did not dominate the analysis, we computed ΔF/F for each cell using the equation:

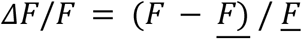

where:

*<u>F</u>* = The mean of *F* across time through the entire session

Cells with very high ΔF/F values (max ΔF/ F > 10), likely not arising from spikes, were discarded from further analysis.

Imaging data was split into individual trials, defined as 2 seconds preceding and 8 seconds following the onset of a trial. Frames occuring while the photostimulation laser was on were excluded due to artifactual crosstalk in the imaging channel (as well as 2 frames prior and after stimulation to ensure there was no contamination from the laser in neighbouring time frames). Trial onset was defined either as the onset of photostimulation in the case of go trials, the onset of galvo spiralling in the case of catch trials or the delivery of reward in the case of reward only trials.

### Definition of targets, photostimulation responsiveness, and excited/inhibited neurons

Neurons were defined as ‘targeted’ on an individual trial if any part of their cell body was located within 15 μm of the centre of the photostimulus beamlet. Neurons were categorised as responsive or unresponsive to stimulus or reward in a trial-wise manner. For each trial, the distribution of ΔF/F values 500 ms pre-stimulus were compared to 500 ms post-stimulus for each cell. A cell was deemed as responsive to a stimulus on an individual trial if it passed a significance threshold of p = 0.05, using a two-sided Wilcoxon signed-rank test, following false discovery rate (FDR) correction. The alpha of the FDR correction (0.015) was set empirically as the value which yielded ∼5% of cells responding on correct rejection trials, where there was no stimulation or licking response. Following significance testing, cells were split into positively and negatively responding cells based on whether the mean ΔF/F value post-stimulus was greater or less than the mean ΔF/F value pre-stimulus respectively. While calcium imaging with GCaMP does not visualise inhibition explicitly, it has been shown that deviations of fluorescence traces below baseline is indicative of inhibition of tonically active neurons^79^. The fraction of neurons excited or inhibited on a given trial, in a given brain region, was defined as the number of positively or negatively responding cells respectively, divided by the total number of neurons recorded in that brain region on that session.

### Pre-stimulus population metrics

All pre-stimulus population metrics were computed across a 500 ms period immediately prior to photostimulation. All metrics were calculated on a trial-wise basis. The natural logarithm was taken of metrics that were fit better by a log-normal distribution as opposed to a normal distribution as assessed by Kullback-Leibler divergence (for clarity, this includes population variance).

Mean population activity was computed by first averaging ΔF/F activity across all pre-stimulus frames for each neuron to yield a vector containing a scalar value for each neuron defining its pre-stimulus activity. Next firing rates were averaged across all neurons to give a single scalar value for each trial, defining the average population activity.

Population variance was computed in a similar fashion, first by averaging across all pre-stimulus frames (in the 500ms window prior to stimulation) for each neuron. However rather than taking the mean of the activity vector as above, the variance of the vector was used to generate a single scalar value for each trial.

### Normalisation and sorting of post-stimulus neural activity

Post-stimulus neural activity was baselined relative to pre-stimulus activity in order to assess the relative change in activity after the photostimulation period, and to compare this change across cells and trials. On each trial, for each individual cell, the average ΔF/F activity in the 2 seconds preceding the photostimulation was subtracted from the post-stimulus activity trace. This normalisation procedure was applied to all analyses and visualisations of post-stimulus neural activity, except for **Fig. 2d-f** and **Extended Data Fig. 5a** where the difference between pre-stimulus and post-stimulus activity was assessed. Neurons were sorted for visual clarity only (in **Fig. 2**), using the sum of the post-stimulus ΔF/F activity on hit, miss and reward only trials. This yields a sorting from strongly inhibited to strongly excited cells.

### Logistic regression classifiers

The dynamic decoding classifiers of Fig. 3 used a logistic regression model with an L2 penalty term to the weights with a regularisation strength of 0.001 (optimised by a parameter sweep with values 10^−7^ to 10^3^ with increments of factor 10). The Scikit-learn implementation of logistic regression was used^119^. Classification accuracy was computed per time point on a session-wise basis and then averaged across sessions, with shaded areas showing the 95% confidence interval of performance across sessions.

Trials of the majority trial type class were randomly subsampled to adjust for sessions in which there was an unequal number of different trial types and ensure that models were not biased to a major or minor class. This meant that the reward only / correct rejection classifiers (**Extended Data Fig. 6e, f**) could only use 10 trials: the number of reward only trials. To ensure that this low number of trials did not underlie the difference in performance with the hit / correct rejection classifier, we also trained a hit / correct rejection classifier using 10 trials only. For this analysis, we sampled 10 hit trials (for each time point) according to the response time distribution of reward only trials, to also control for the difference in response times between the two trial types (**Extended Data Fig. 2d**). We found that hit predictions were significantly greater than reward only predictions on all post-stimulus time points in S1 and 56/110 post-stimulus time points in S2 (**Extended Data Fig. 6g, h**).

Each model was trained to classify the probability that a trial belonged to one of two different trial types. A 3:1 train:test split was employed and model performance was assessed on held-out test trials only. 4-fold cross validation was used on each session, with a new model trained for each fold for each time point, and classification accuracy is reported as the average of the test data across folds, meaning all (potentially subsampled) trials were in the the held-out test set exactly once. A new model was trained from scratch for each imaging frame within a trial, hence the training data consisted of a vector containing a single scalar *ΔF/F* value for each cell of all trials on a given frame.

Significance of decoding predictions was assessed by two-sided Wilcoxon signed-rank tests, where the predicted classification was compared to chance level (= 0.5). Predictions were binned per 2 time points, such that each test consisted of 22 (2 time points x 11 sessions) paired samples (prediction and the constant chance level). Predictions were said to be significant if their p value was below 0.05, Bonferroni corrected for the number of tests performed per trial type (83, both the pre-stimulus and post-stimulus activity time points, excluding the photostimulation artifact). The binning of 2 time points was used because single time point statistics had too little data to be significant given the strong Bonferroni correction. Hence we chose the smallest bin size (2) through which it was mathematically possible to yield significant p values. Significant time points are indicated in **Fig. 3** and **Extended Data Fig. 6a-h,7** by thick lines, coloured by the trial type they refer to, at the top of each panel. To quantify the difference between predictions of two trial types, the same tests were performed comparing the two trial type predictions directly.

As reward signals in the brain are complex, and depend on whether the reward is expected or unexpected^120^, we confirmed that the reward signal on hit trials and reward only trials was comparable, thus ensuring that the hit vs correct rejection classifier was indeed classifying the neural signature of perceived stimulation and not just a reward signal that is different from the reward only trials **(Fig. 3b, c)**. This is critical as the water reward on hit trials was expected, but not expected on reward only trials. We addressed this by training a model to classify reward only trials from correct rejection trials, thus this classifier should detect only the neural signal from rewards, and related motor activity **(Extended Data Fig. 6e,f)**. Testing this model on hit trials resulted in identical performance to the reward only condition both in S1 and S2 (no significant difference at any time point in S1 or S2). Hence, classifiers trained specifically to detect rewards could not distinguish hit and reward only trials, meaning that the neural response to reward is comparable in both trial types and the classification of hit trials from correct rejection trials is indeed decoding the neural signature of perceived stimulation, both in S1 and S2.

We further verified that the dynamic decoding results of **Fig. 3** were reproduced when larger time windows of 2 seconds were used **(Extended Data Fig. 6i-l)**. Additionally, we verified that the dynamic rise in S2 Hit/CR decoding accuracy during the response window (0ms - 1000ms) of **Fig. 3c** was not solely due to an increase of the number of trials that responded to the stimulation with time (**Extended Data Fig. 2c,d)**. To do so, we considered the first decoder post-stimulus (at t=367 ms), and compared the decoded prediction of individual hit trials (of withheld test data only, eventually evaluating all trials using the previously described cross-validation procedure) with their response times **(Extended Data Fig. 6m)**. We found that no sessions exhibited a significant correlation **(Extended Data Fig. 6n)**, with non-significant correlations spreading across both positive and negative values, showing that there was no consistent correlation between response time and decoding accuracy across sessions.

Next, we assessed whether the stimulus encoding strength depended on the strength of the stimulation (i.e. number of cells targeted) or the S1 population variance, as both these metrics influence the probability of animals to detect the photostimulation (**Fig. 6**). First, we observed that the number of cells targeted led to a significantly different decoding performance directly post-stimulus in S1 only (**Extended Data Fig. 13;** On hit vs correct rejection classifiers, for the 18 time points up to 1s: hit 40/50 was significantly different from hit 5/10 in S1 for all time points; miss 40/50 was significantly different from miss 5/10 in S1 for 16 time points; there were no significantly different time points for the two equivalent comparisons in S2; Wilcoxon signed-rank test, Bonferroni corrected). In S1, stronger stimulation led to more positive decoding performance of both hit and miss trials **(Extended Data Fig. 13)**. This likely indicates that the photostimulation-induced perturbation of cells in S1 was picked up by the classifiers, but that the ensuing network response (in S2 and later in S1) was equivalent for hit and miss trials irrespective of the strength of stimulation. Second, we found that S1 population variance did not influence the stimulus encoding strength in either S1 or S2, except for a slight difference in S1 for miss trials on the hit vs correct rejection classifier (**Extended Data Fig. 9**; On hit vs correct rejection classifiers, for the 18 time points up to 1s: miss high-population-variance was significantly different from miss low-population-variance in S1 on 8 time points; there were no significantly different time points for the equivalent tests in S2 and for hit trials in both S1 and S2; Wilcoxon signed-rank test, Bonferroni corrected).

### Behavioural data analysis

As imaging was stopped intermittently, neural activity was not recorded for every trial performed by the animal; trials that were not imaged were excluded from all analysis. Hit trials in which the animal licked with an exceptionally short-latency (<150 ms) are likely to have been driven by random licking rather than perception of the stimulus and were thus marked as ‘too-soon’ and not included in further analysis^32^.

For trial-wise analysis of neural activity, each trial’s time series was aligned to the onset of the (sham) photostimulation (time = 0s) or to the onset of reward for the reward only condition.

Psychometric curves were fit to behavioural data by computing the value of d’ separately for trials in which a given number of cells was targeted (**Fig. 1f**). This was achieved by comparing the hit rate for a given number of cells targeted to the false positive rate across all catch trials. Data was fit using a logistic function adjusted lower bound at d’ = 0.

### Quantification of SNR effect

We performed a logistic regression analysis to validate that both stimulus strength (i.e., the number of targeted cells) and the noise level (i.e., pre-stimulus population variance) significantly contributed to predicting trial outcome (**Fig. 6**). Logistic regression was performed on the data of all hit and miss trials of all sessions (as in **Fig. 6a,b**). Both regressors significantly contributed to explaining trial outcome (both p values < 10^−13^, two-sided t-test on the logistic regression coefficients), with explained variance R^2^ of 8.4% (as calculated by McFadden’s pseudo-R^2^ for logistic regression). This was greater than the R^2^ obtained by regressing trial outcome to each of the two variables individually (R^2^ = 5.4% for number of targeted cells and R^2^ = 2.7% for population variance). Hence, trial outcome was best predicted using both regressors.

### Calculating the across-neuron population-wise correlation

We calculate the across-neuron population-wise correlation before photostimulation onset as an indicator of coordination between neurons, similar to that described in^48^. The population-wise correlation *ν* is defined as the variance explained by the first latent factor of neural activity across trials of the same condition. To calculate correlations as precisely as possible but avoid any bias from different trial numbers between conditions and sessions, we concatenate 6.5 seconds of pre-stimulation activity from 15 sub-sampled trials at a time, and report the average of population-wise correlation *ν* across 1000 sub-samples for each session and condition.

### Calculating the effective recurrency from the covariance matrix of non-shared activity

We infer the effective recurrence (*R*) from the covariance matrix of non-shared activity as described in^57^. As a rough outline, the three main steps are (1) to disentangle the activity that is shared across all neurons from the non-shared activity that is individual to each neuron. (2) From the non-shared activity, estimate the width of the cross-covariance distribution, which is an indicator of the dynamical state of the system, and the mean variance. (3) From the ratio of these two quantities, calculate the effective recurrence R directly as derived in^52^.

#### Preprocessing

For the subsequent analysis, we want to identify whether the recurrence *R* is predictive for the detection of cortical stimulation. Hence, for the analysis we need a sufficiently long period of stationary activity preceding stimulus onset. We therefore used the window from -7 s to -0.5 s prior to stimulation **(Extended Data Fig. 10a,b)**. To obtain a bin size equal to the 100 ms bins used in Dahmen et al.^57^, we re-binned the baselined ΔF/F, which was recorded with a framerate of 30 Hz (i.e. one frame is 33 ms long), by averaging over 3 adjacent frames. This yields 65 bins for each pre-trial interval.

#### Latent Factor Analysis

We separate correlated activity, which might present common external input, from non-shared activity, using Latent Factor Analysis (LFA)^57^. The LFA was applied to the activity matrix created by concatenating all of the pre-hit and pre-miss intervals in a single session, yielding a matrix of size *n*_*neurons*_ × (65 × *n*_*trials*_) **(Extended Data Fig. 10d)**. We discarded the activity explained by the first 5 latent factors and used the non-shared activity that is not explained by the first 5 factors **(Extended Data Fig. 10f)** for subsequent analysis.

In more detail, Latent Factor Analysis (LFA) represents the observed activity *X* (minus its mean activity *M*) as a linear combination of 5 latent factors – each weighted with a loading matrix *L* - plus a non-shared term *E* that is different for each neuron:

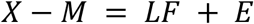

The choice of 5 latent factors for *n*_*neurons*_ = 357 ± 116 was made for all sessions in parallel to ensure consistency across sessions, and closely follows the criterion introduced in ^57^: We calculated the min-max normalized 5-fold cross-validated log-likelihood of the LFA fit for each session, and plotted its increase per factor **(Extended Data Fig. 10d)**. We now chose the maximum number of factors that shows more than 5% increase in normalized log-likelihood - which is 5 in our dataset, similar to the values reported in Dahmen et al.^57^ (2-10 factors for 200-300 neurons). We also checked the stability of our results with respect to this parameter, and found that they are qualitatively similar for 2-10 factors **(Extended Data Fig. 12a-j)**. In addition, we also confirmed that using Principal Composition Analysis (PCA) instead of LFA yielded equivalent results (**Extended Data Fig. 12k,l)**. Latent Factor Analysis was performed using the FactorAnalysis function from sklearn^119^, which implements a maximum-likelihood estimate of the loading matrix *L* using a randomised and truncated singular value decomposition of *X* − *M*, as is the default. Principal Component Analysis was performed using the PCA function from sklearn^119^, which uses the LAPACK implementation of randomized truncated Singular value decomposition to find the loading matrix of *X* − *M*.

#### Two-Step Bias Correction

The effective recurrency *R* is calculated from the moments of the entry distribution *P*(*c*_*ij*_) of the sample covariance matrix *C* **(Extended Data Fig. 10g)** of the non-shared activity *E*. Since the second moment (the variance) of this entry distribution is biased by the number of trials used to calculate the covariance matrix, and the number of trials differs between hit and miss conditions (*n*_*hits*_ = 92 ± 37, *n*_*misses*_ = 48 ± 36, *min*(*n*_*hits*_, *n*_*misses*_) = 16), we performed a two-step re-sampling procedure to avoid this bias.

#### Outside step

In the outside step, we first choose a random sub-sample of 15 hit or miss trials without replacement to ensure that the subsequent analysis steps are performed with the same number of trials. This outside step is performed for either all 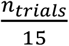 possible permutations of 15 trials if the number of permutations is less than 1000, or else 1000 randomly chosen permutations. Using these 15 trials, we calculate the moments of the entry distribution *P*(*c*_*ij*_) of the non-shared covariance matrix. The mean on-diagonal covariance *µ*_*V*_ is unbiased with respect to the number of bins *T* used to calculate the covariance matrix, and can therefore be calculated directly using all bins from 15 trials:

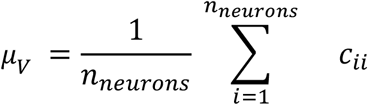

#### Inside step

The standard deviation of the off-diagonal covariance is over-estimated if only a small number of bins *T* is used. To correct for this, we performed the inside step where we extrapolate to *T →* ∞. This is done by sub-sampling the number of bins *T* even further and using the expected analytical relation between the over-estimate 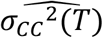and the sub-sampled number of bins *T* as derived in^57^: In detail, the asymptote is obtained by fitting the relation

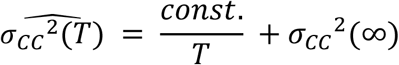

to the biased estimate of the standard deviation calculated from *T* bins **(Extended Data Fig. 10j)**. This fit is done by applying the curve_fit function from scipy^112^ to the average of 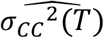for 50 repetitions of choosing *T* = [5 ∗ 65,6 ∗ 65,. ‥, 15 ∗ 65] bins without replacement (but respecting trial identity) from the 15 trials selected in the outside step – this fit completes the inside step.

#### Final Estimate

To complete the outside step, an estimate of *R* is calculated from the ratio of *σ*_*CC*_ ^2^(∞) and *µ*_*V*_^2^ as

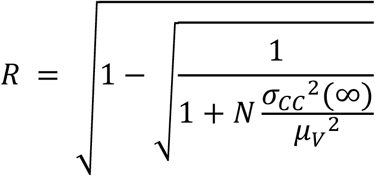

where the free parameter *N* = 50000 is chosen to match the number of neurons in the immediate vicinity of the field of view^57^. Finally, the estimates of *R* obtained for each outside step are averaged across all combinations to obtain an *R* number for each session and condition. Note that the two-step resampling procedure is computationally intensive: performing the parameter sweep shown in **(Extended Data Fig. 12)** takes multiple days on ∼25,000 CPU cores.

#### Correlation

Mean pairwise correlation was computed on the concatenated activity after performing the Latent Factor Analysis by calculating the cross-correlation matrix for all pairs of neurons, whereby element i,j in this matrix was the Pearson correlation between neuron i and neuron j across all pre-stimulus frames. A single scalar value was generated for each trial type by computing the mean across all off-diagonal elements of this matrix.

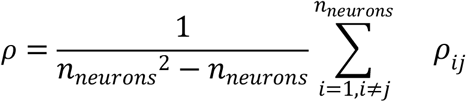

### Calculating the network response timescale from photostimulated neurons

We calculated the network response decay time after photostimulation as a measure of persistent activity likely generated by local recurrence, adapted from^59^. To capture as much of the response as possible, we restricted our analysis to the subset of trials where photostimulation lasted 250 ms (i.e. 5-50 cells targeted). For each trial, we selected only the targeted neurons and averaged their photostimulation response up to 5.5 s corrected for their average fluorescence in a 6.5 s pre-trial window. We then computed the average response across trials, sub-sampling 10 trials at a time 1000 times to avoid any bias from different trial numbers between conditions and sessions. Last, we fit an exponential decay function with amplitude *A* and decay time *τ*_*post*_ to the averaged responses, and finally reported the average of the decay time *τ* across 1000 sub-samples for each session and condition. To relate the network response timescale *τ*_*post*_ to the effective recurrence *R*, we employed a simple linear network model^121^ which, simply put, describes the firing rates of the targeted neurons **r**(*t*) as a function of the recurrent weights **W** and external input **h**(*t*). After an eigenvector decomposition of **W** and identifying the largest eigenvalue as *R*, we find the decay timescale along its eigenvector *τ*_*post*_ to scale with *R* as 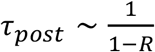.

### Statistical procedures

We did not use statistical methods to determine sample size and no randomisation or blinding was used. Unless otherwise stated, paired non-parametric tests were employed and a p value of 0.05 was used as a threshold for significance throughout. Multiple comparisons corrections were applied to significance tests using the Bonferroni correction unless otherwise stated. Error bars show 95% confidence interval unless otherwise stated.

## Author contributions

J.M.R. and A.M.P. designed the study. J.M.R and T.A. developed the behavioural control apparatus and software. J.M.R., R.M.L., and J.K. conducted experiments. J.M.R., T.L.v.d.P., and M.L., performed analysis with advice from J.D., V.P., and A.M.P‥ J.M.R., T.L.v.d.P., M.L., and A.M.P. wrote the paper with input from all co-authors.

## Acknowledgements

We thank Andrew J. King, Armin Lak, Simon Butt, Andrew Saxe, Christopher Summerfield and Christopher D. Harvey for useful discussion and Bruker Corporation for technical support. This work was supported by funding from the Wellcome Trust (204651/Z/16/Z) to J.M.R., R.M.L., and A.M.P., from the Biotechnology and Biological Sciences Research Council grant number BB/M011224/1) to T.L.v.d.P., from the Max-Planck Society to M.L. and J.D‥, from SMARTSTART to M.L. and J.D., from the Erasmus mobility program to M.L, and from the German Research Foundation (DFG) to VP (SPP 2205, Project Number 430157073).

## Extended data

**Extended Data Figure 1:**
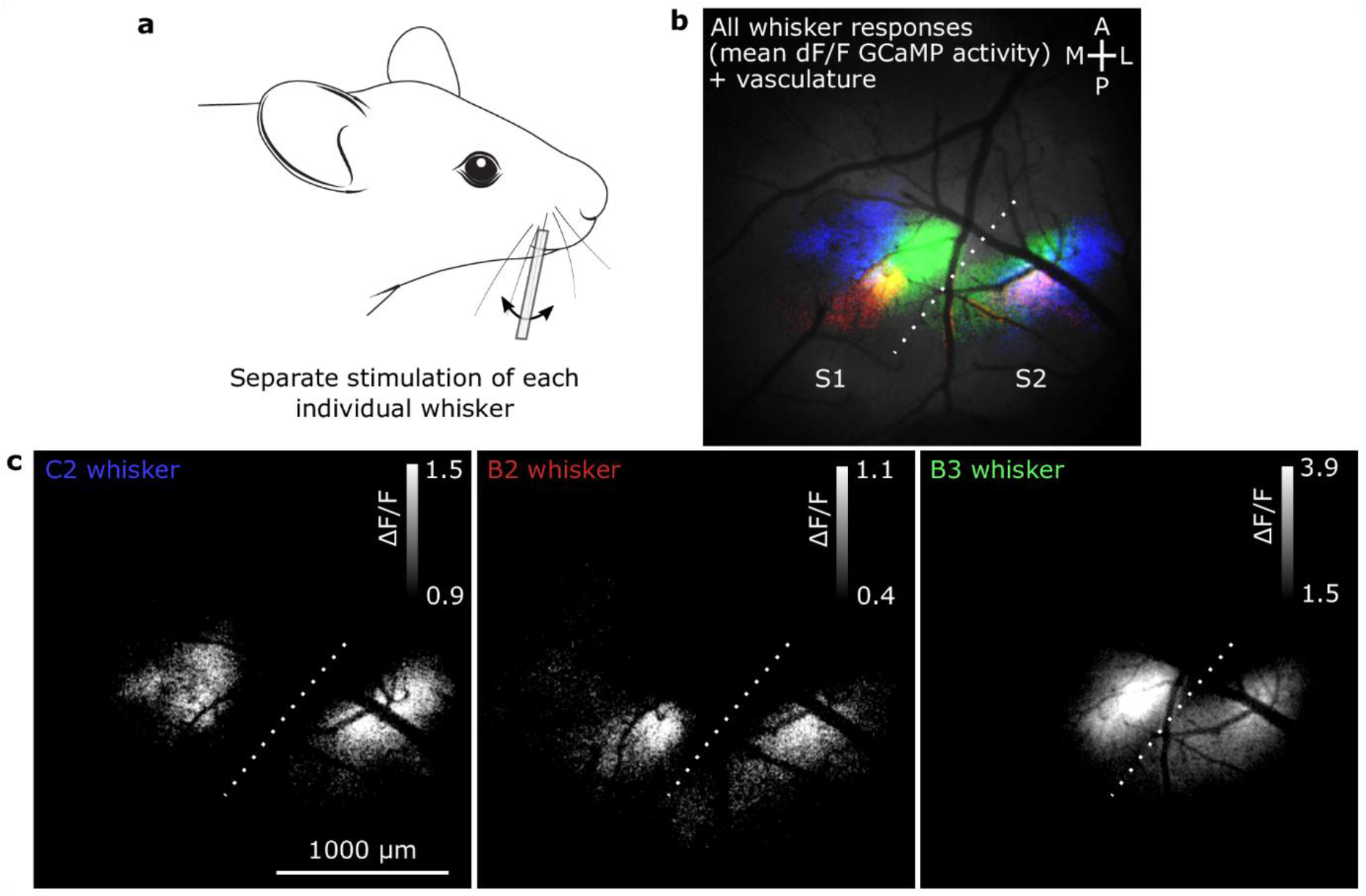
Mapping whisker S1 and S2 regions using widefield calcium imaging and whisker stimulation. Single whiskers were deflected one at a time by threading them on to a capillary tube attached to a piezoelectric actuator. Each whisker was deflected multiple times and the results were averaged. The stimulus-triggered average responses from all whisker deflections is plotted on an image of the cerebral vasculature. **(c)** Individual stimulus-triggered average images from each whisker shows the topographical organisation of the barrel cortex in whisker S1 and the mirrored topography in whisker S2. The drawing in **(a)** was adapted from a drawing by Ethan Tyler and Lex Kravitz (Scidraw.io, doi.org/10.5281/zenodo.3925901).

**Extended Data Figure 2:**
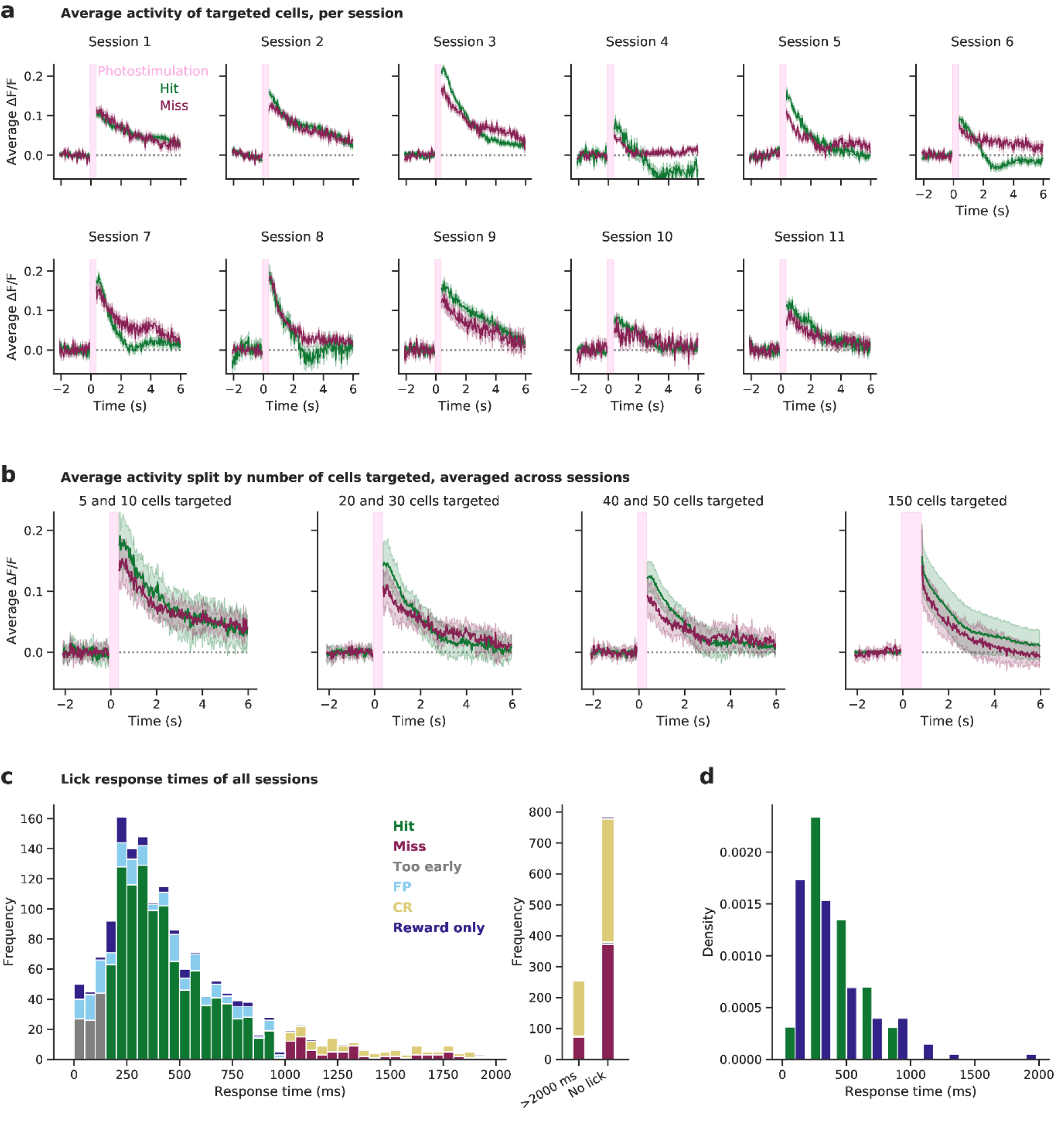
Activation of targeted cells by photostimulation. **a)** The average ΔF/F activity of the cells targeted by photostimulation is shown per recording session, for Hit and Miss trials separately. Targeted cells are activated similarly, with the notable exception of Hit-only inhibition at ∼2 seconds post-stimulus in 3 recording sessions (of the same animal), indicating that the photostimulation-induced activation is largely independent of behavioural outcome. All trials with 5 to 50 cells targeted were used; trials where 150 cells were targeted were excluded because almost all of these trials resulted in a Hit outcome, biasing the averages. **b)** The same data as panel a) is shown, but now averaged across animals and split by number of cells targeted. **c)** Stacked histogram of the animals’ response times (defined by the time of the first lick) for each trial type. All trials with a response time within 2000 ms of all recording sessions are shown in the left panel, while trials with a response time greater than 2000 ms and trials where no lick occurred are grouped in the right panel. **d)** Density histograms of response times of hit and reward only trials. Same data as in panel a), but normalised per trial type and binned per 200ms. Medians (248ms for reward only trials and 383 ms for hit trials) are significantly different (p=0.01, Mood’s median test).

**Extended Data Figure 3:**
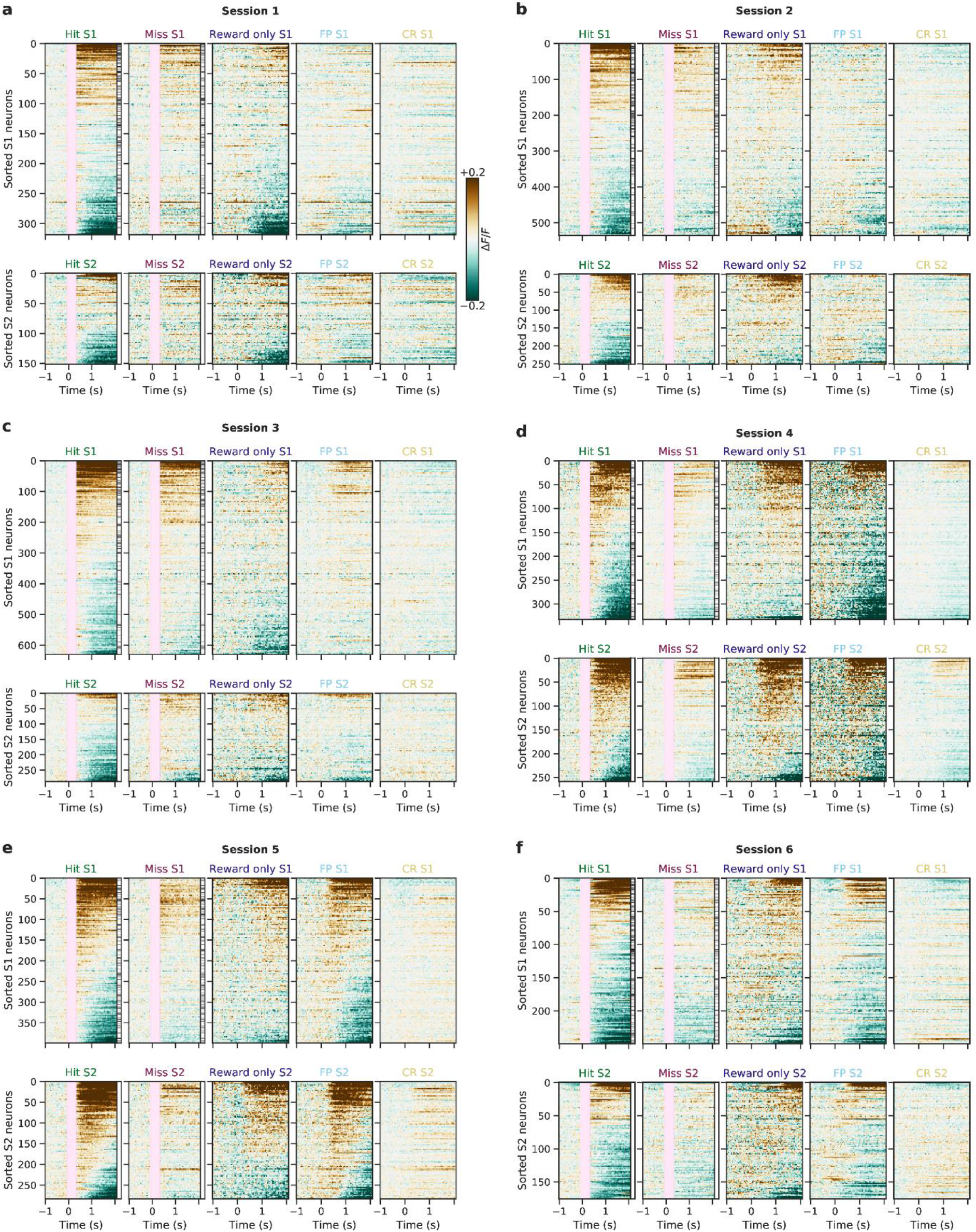
Trial-averaged raster plots of all sessions and trial types. **(a-f)** Raster plots showing the trial-averaged responses for all trial types to photostimulation (pink vertical bar, hit/miss) and/or reward (hit/reward only) and/or licks (hit/reward only/false positive) and none of these (correct rejection) of individual cells from the first 6 sessions.

**Extended Data Figure 4:**
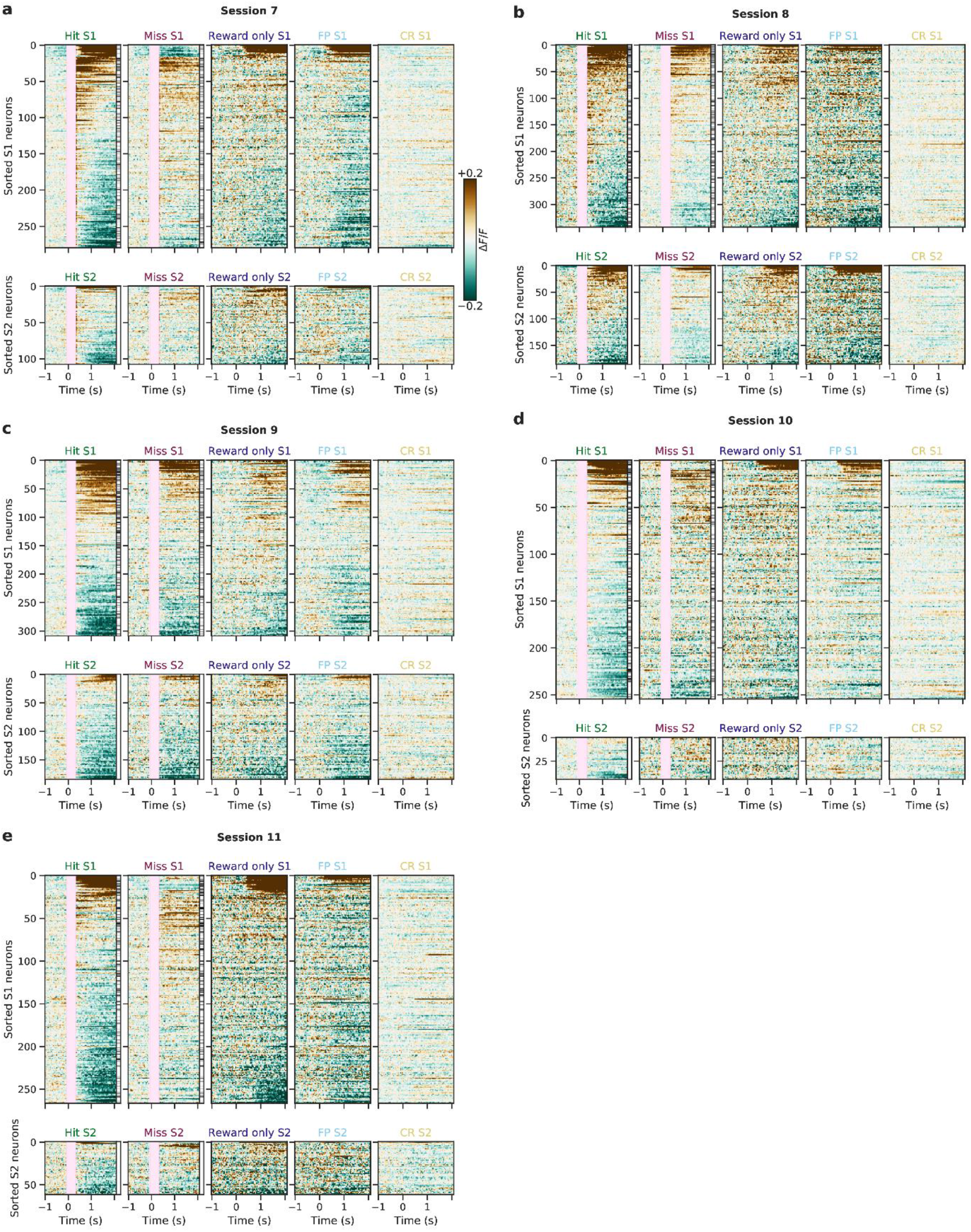
Trial-averaged raster plots of all sessions and trial types: continued. **(a-e)** As above for the remaining 5 sessions.

**Extended Data Figure 5:**
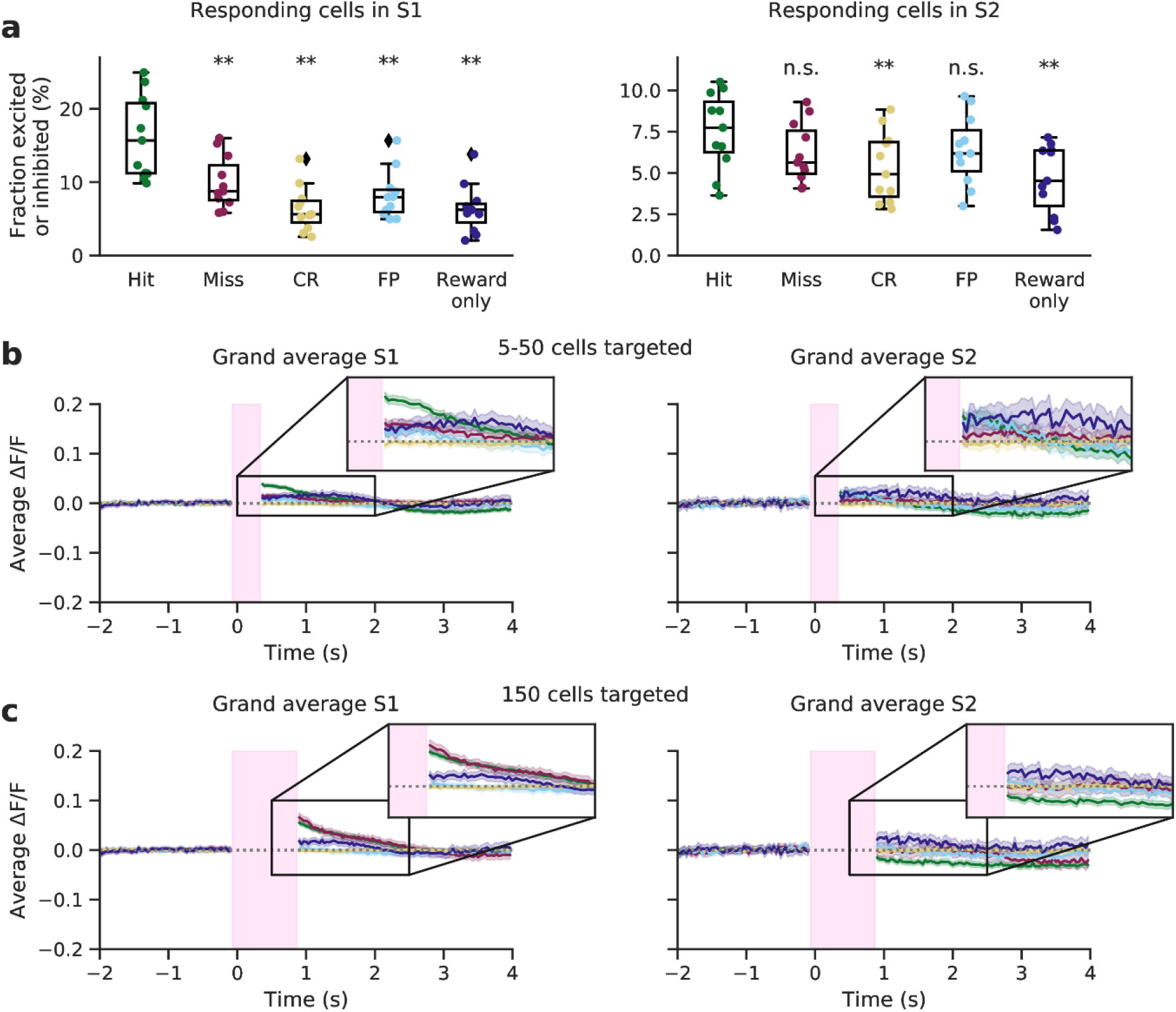
Fraction of cells responding and grand average traces of all trial types in S1 and S2. **(a)** The fraction of responsive cells (both excited and inhibited) on each trial type across all numbers of cells targeted. Wilcoxon signed-rank tests were used to test whether the fraction of responsive cells of hit trials was significantly greater than on other trial types (** indicates p values < 0.05, Bonferroni corrected for 4 tests). **(b)** Average population responses of all trial types across all sessions. Traces are averaged across cells, trials and sessions for a given trial type. Trials in which 150 cells were targeted were removed for display (and shown in **c**). The time course of reward only trials is different from the time course of hit trials, hinting that the neural activity on hit trials constitutes more than motor preparation, movement, and reward related activity. This is further quantified in **Fig. 2, 3. (c)** As above but showing exclusively trials in which 150 cells were targeted.

**Extended Data Figure 6:**
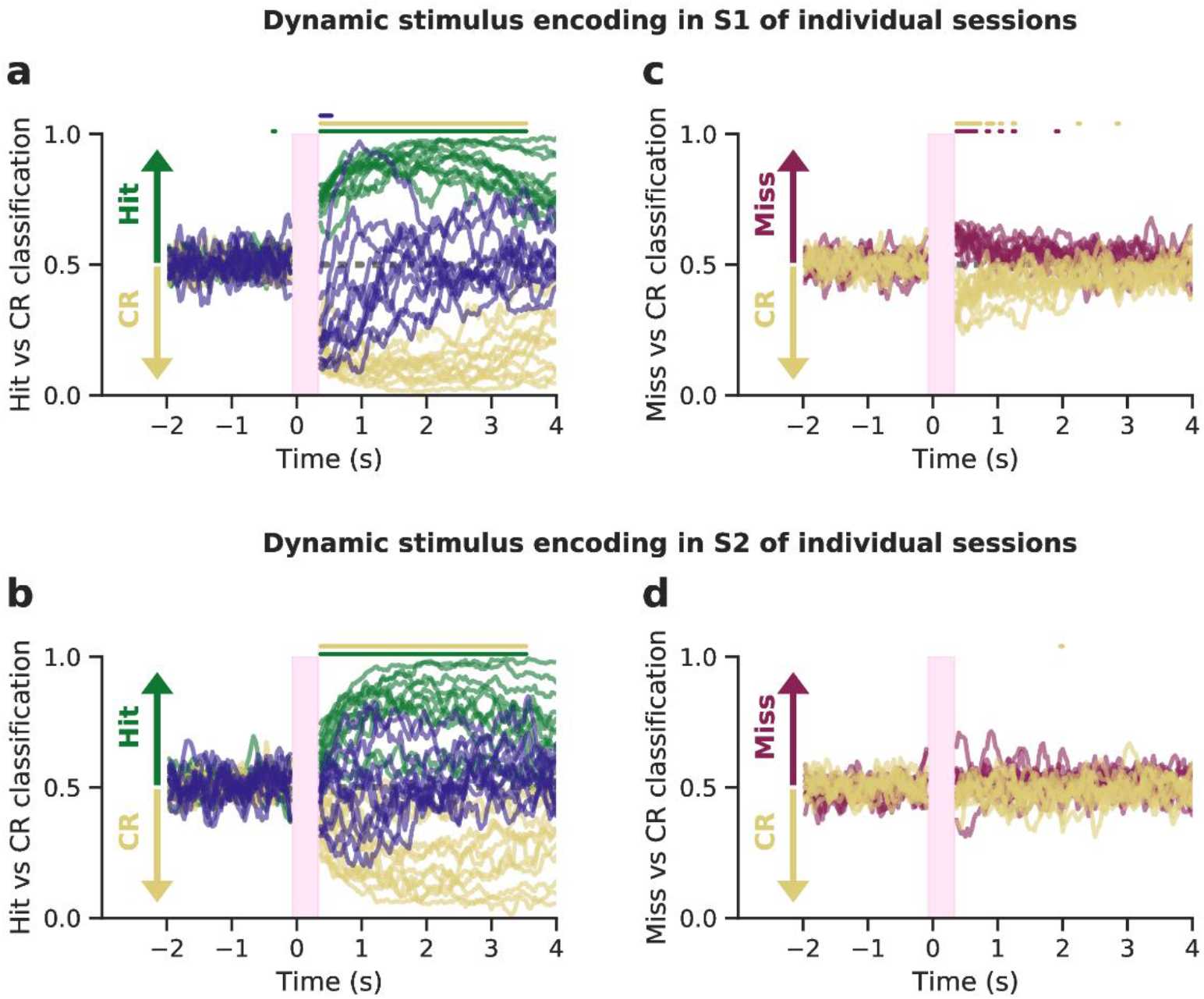
Dynamic decoders of all trial types. The strength of stimulus decoding of trial type in the neural population in S1 was dynamically quantified using logistic regression models. Models were trained on each time frame individually, with activity of all cells in S1 or S2, and tested on held-out data. **(a)** Models were trained, for each time point, on S1 activity to classify hit trials from correct rejection trials and then tested on held-out hit trials (green), correct rejection trials (yellow), reward only trials (dark blue), false positive trials (light blue) and miss trials (red). Coloured bars above the traces show time points at which classifier performance was significantly different from chance (p < 0.05, Bonferroni corrected, see **methods**). **(b)** As above, but trained on S2 activity. **(c)** Models were trained on S1 activity to classify miss trials from correct rejection trials and then tested on miss trials (red), correct rejection trials (yellow) reward only trials (dark blue), false positive trials (light blue) and hit trials (green). (**d)** As above, but trained on S2 activity. **(e)** Models were trained on S1 activity to classify reward only trials from correct rejection trials and then tested on reward only (dark blue), correct rejection trials (yellow), miss trials (red), false positive trials (light blue) and hit trials (green). **(f)** As above, but trained on S2 activity. **(g)** As (a) but the number of trials was restricted to 10 for each type, matching the total number of reward only trials recorded. (**h)** As (b) but the number of trials was restricted to 10 for each type, matching the total number of reward only trials recorded. **(i-l)** Equivalent to Figure 3, but now using 2-second windows to train the classifiers. Three windows were used (-2.0s up to and including -0.1s, 0.4s u/i 2.3s, 2.4s u/i 4.3s), shown at the bottom of each panel. For each window, neural activity was averaged across time, per neuron per trial. Classifiers were then trained and evaluated as before and described in Methods. Asterisks indicate significant differences with chance level accuracy 0.5 (Wilcoxon signed-rank test, bonferroni-corrected p value < 0.05). **(m)** Comparison between lick response time and predicted outcome for all hit trials in S2. We considered the first decoder post-stimulus (at t=367 ms), and compared for each trial the predicted outcome evaluated on withheld test datato the response time. This panel shows one example session (same session as Fig. 2a), and its Pearson correlation value r and associated (two-sided) p value. **(n)** Pearson correlation values r between response time and decoded S2 predictions of hit trials of all 11 sessions are shown. A Bonferroni multiple-comparison correction of N=11 (sessions) was used.

**Extended Data Figure 7:**
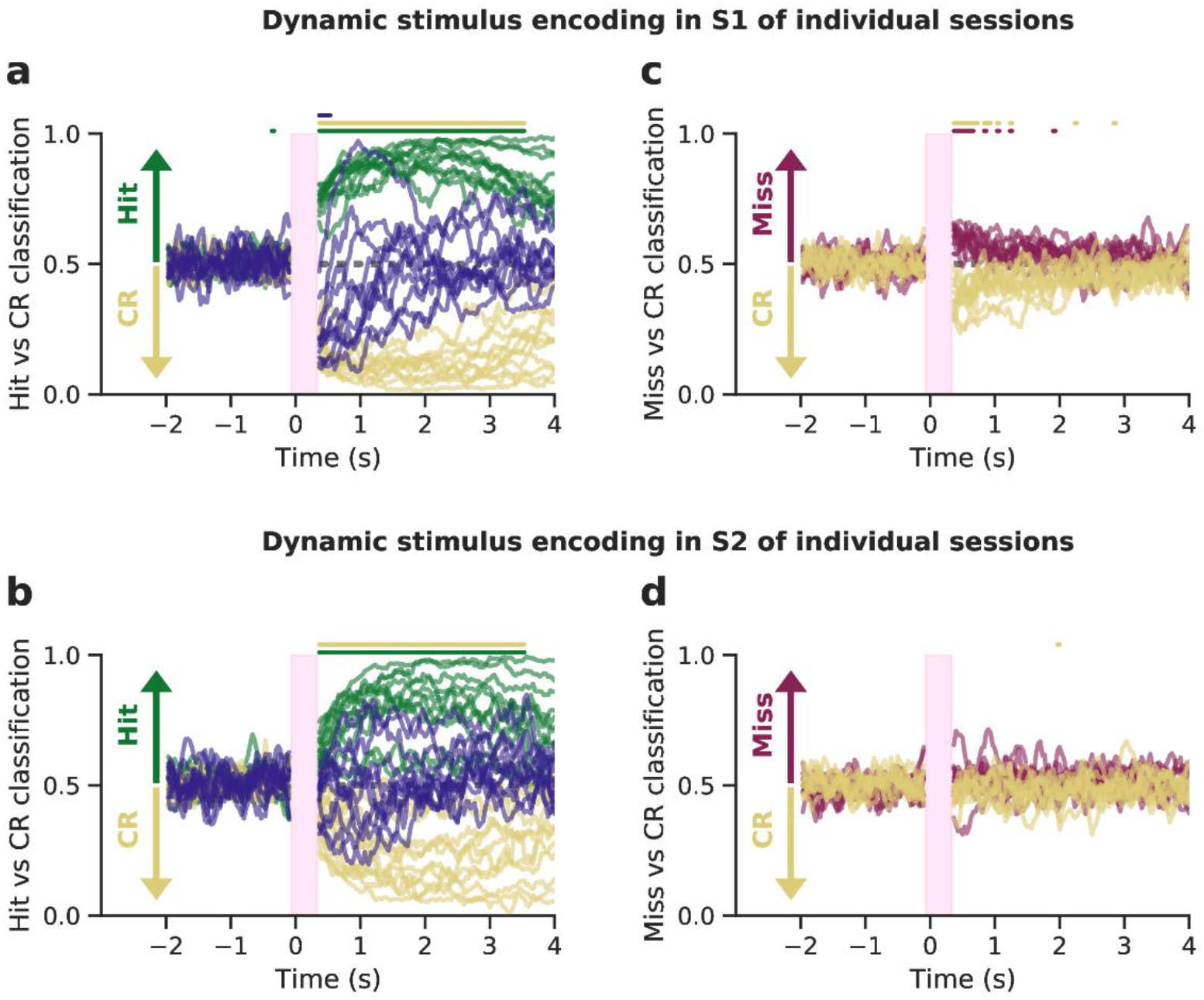
Dynamic decoders of individual sessions. As **Fig. 3**, but displaying classifier performance for each individual session. Traces were smoothed using a running mean of 5 time points for visual clarity.

**Extended Data Figure 8:**
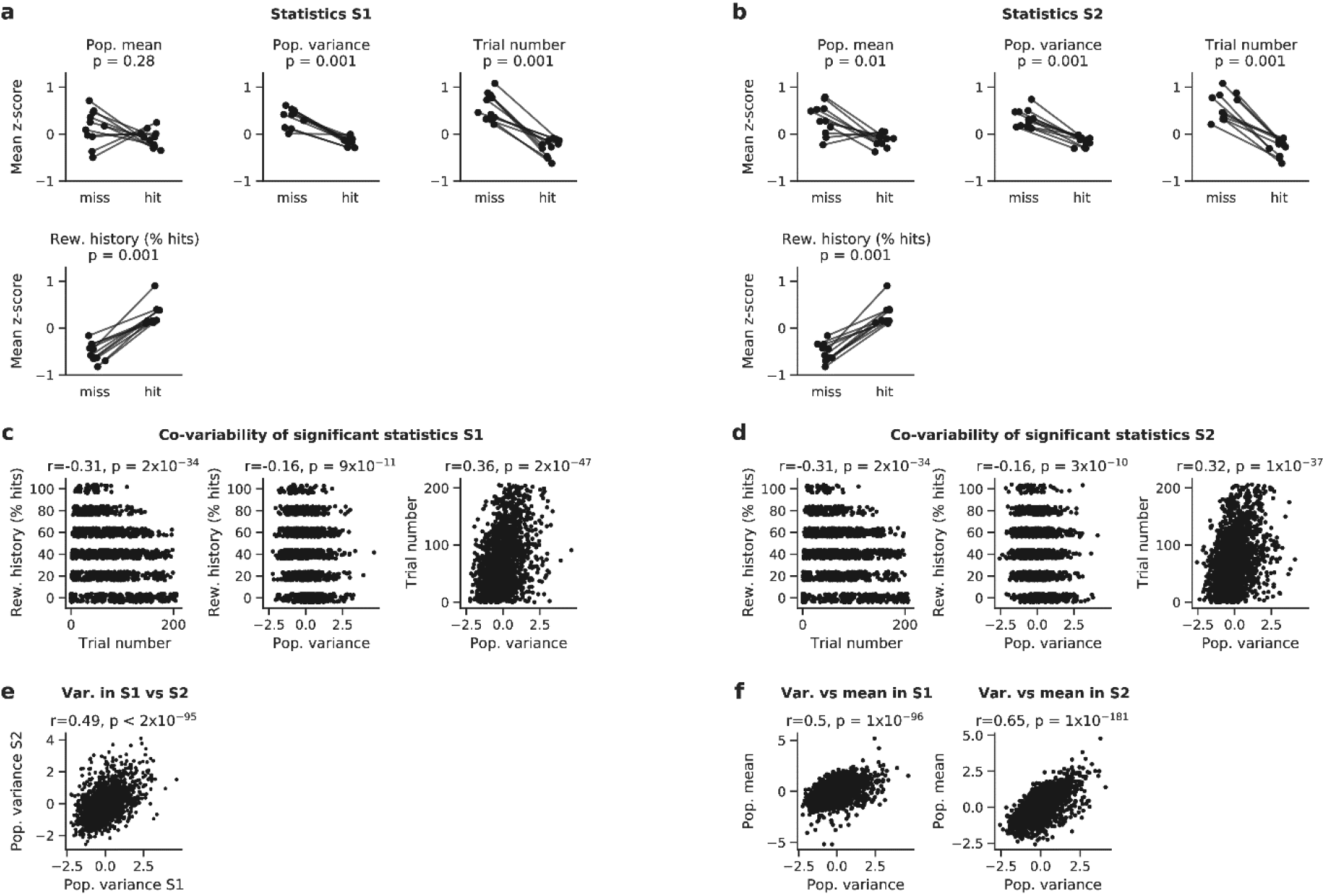
Further population metrics of pre-stimulus activity. Comparison of population metrics of pre-stimulus in S1 (**a)** and S2 (**b**) activity prior to hit trials and prior to miss trials. The first two panels show the two metrics of **Fig. 4b**. Next, two other significant task variables are shown; the trial number (an integer between 0 and the total number of trials in a session, indicating the number of trials previously undertaken by the animal in a given session) and the reward history (defined as % hit trials in the 5 preceding photostimulated trials). All variables were z-scored for clarity, and significance was assessed using Wilcoxon signed-rank tests. (**c, d**) The co-variability of the significant statistics of panels **a** and **b** was assessed using Pearson correlation. In sum, as recording sessions progress, reward history declines, and population variance is correlated to this trend. **(e)** Population variance is significantly correlated between S1 and S2 on a single-trial level. **(f)** Population variance is significantly correlated to the population mean on a trial-by-trial basis in both S1 and S2. Population mean and variance were z-scored in all panels to allow comparison between different recording sessions. Trial number and reward history were z-scored in panels **a-b** only.

**Extended Data Figure 9:**
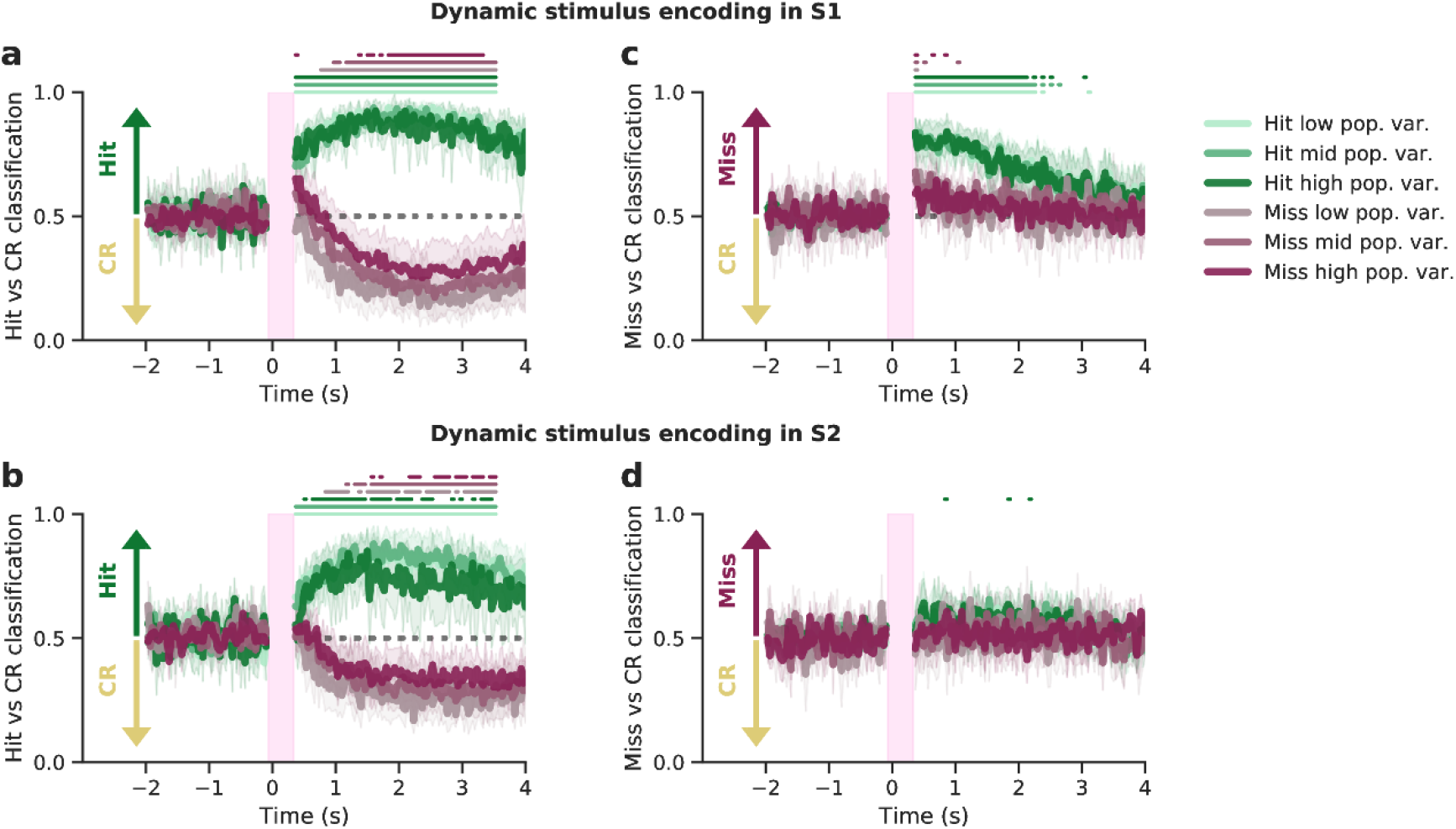
Dynamic decoding results split by population variance. The decoders of **Fig. 3** are shown, but additionally split by population variance. Each recording session was split into 3 tertiles of equal size based on population variance (i.e. 3 bins were used), which were then averaged across sessions.

**Extended Data Figure 10:**
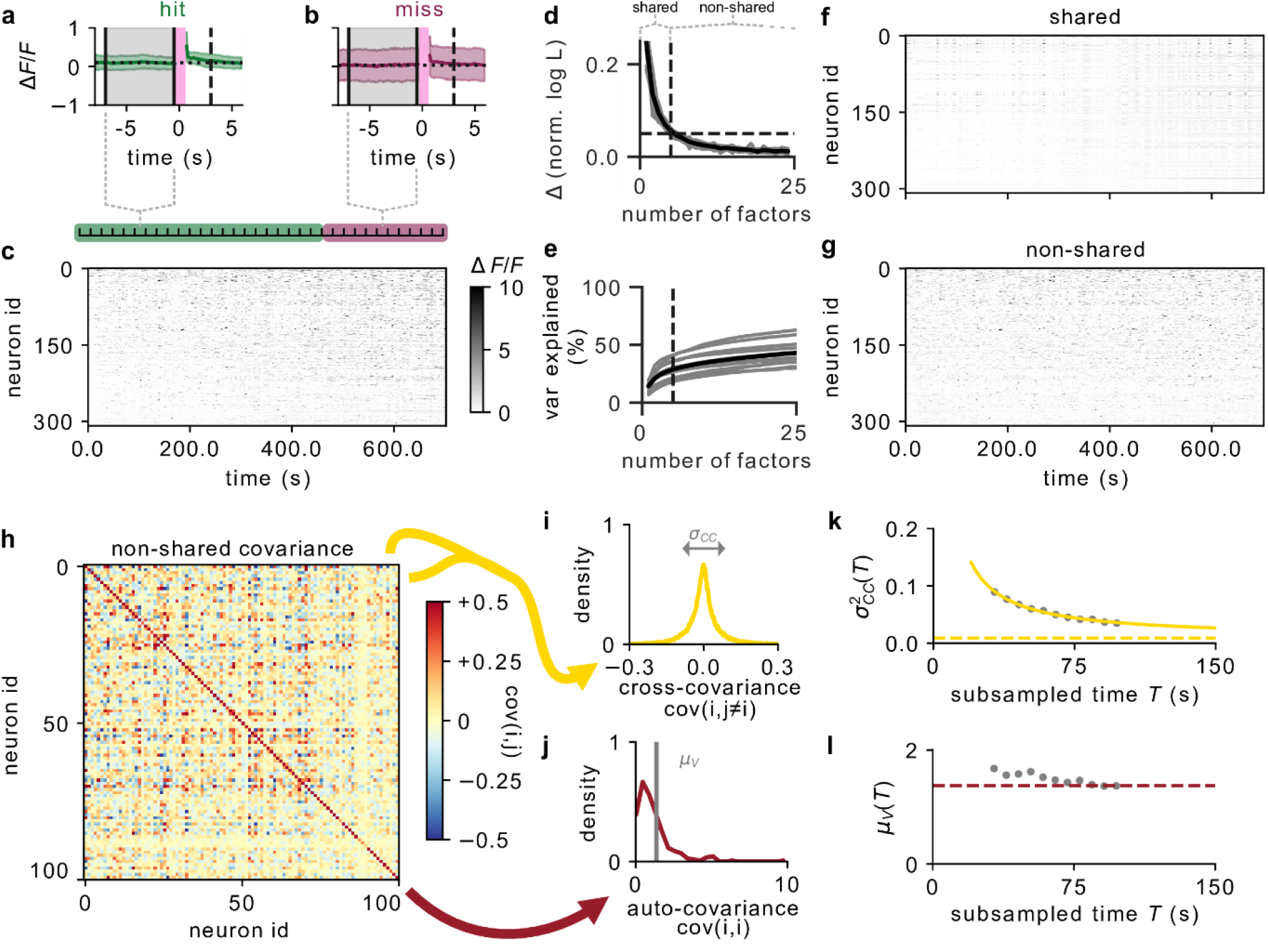
Inferring the effective recurrence from non-shared covariances. **(a)** Trial-averaged peri-stimulus average activity of S1 during hit trials in an example session (Mouse RL123, run 22, 87 hit trials). The green line and shaded region show the mean ΔF/F and variance across all S1 neurons and all hit trials. The two vertical solid black lines are the bounds of the (pre-stimulus) analysis period, which is coloured in grey. The single vertical dashed black line also denotes the start of the analysis period, but here referenced to the preceding stimulation (the earliest possible start, as the inter-trial-interval varies). The horizontal dotted black line denotes the mean population activity during the pre-hit period. The pink bar denotes the photostimulation period. **(b)** Trial-averaged peri-stimulus average activity of S1 during miss trials (red). **(c)** Pre-stimulation activity of S1 neurons concatenated across all 87 hit and 21 miss trials, trial structure shown at the top is for visualisation only and not to scale. **(d)** Increase in log-likelihood of the LFA fit (min-max normalized) as a function of the number of latent factors performed across all analysis windows preceding hit and miss trials. Grey lines show individual sessions, the black line shows the mean across sessions. The dashed horizontal line shows the 5% threshold that was used to determine the number of factors, the dashed vertical line shows the 5 factors we selected to calculate the shared activity, all other factors were discarded and the non-shared activity is used for further analysis. **(e)** Percentage of the variance explained by the LFA as a function of the number of latent factors. **(f)** Shared pre-stimulation activity of S1 neurons concatenated across all hit and miss trials, as calculated by projecting the first 5 latent factors back onto the neurons. Colour scale is the same as in **c. (g)** Non-shared pre-stimulation activity of S1 neurons concatenated across all hit and miss trials, as calculated by subtracting the shared activity. Colour scale is the same as in **c. (h)** Covariance matrix of non-shared activity in S1, i.e. the activity that is not common across most recorded neurons, that is, not explained by the main 5 latent factors. Only 100 out of 309 neurons are shown. **(i)** Distribution of off-diagonal components *c*_*ij,i* ≠ j_ of the non-shared covariance matrix shown in (a). The grey arrow denotes the standard deviation *σ*_*CC*_ of the on-diagonal components. **(j)** Distribution of on-diagonal components *c*_*ii*_ of the non-shared covariance matrix shown in (a). The grey line denotes the mean *µ*_*V*_ of the on-diagonal components. From the ratio of *σ*_*CC*_ and *µ*_*V*_, the effective recurrence *R* can be inferred. **(k)** Artificial sub-sampling procedure to obtain an estimate of the standard deviation *σ*_*CC*_ of the off-diagonal covariances that is not biased by the finite number of bins *T* recorded in a single session. Grey points show the average of 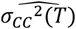for 50 random samples of *T* = [5 ∗ 65,6 ∗ 65,…, 15 ∗ 65] bins chosen without replacement, yellow line shows the fit of the Relation 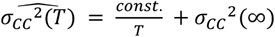, and the dashed yellow line shows the asymptote *σ*_*CC*_^2^(∞) obtained from said fit. **(l)** The mean of the on-diagonal covariances is not significantly biased by the finite number of bins *T*, hence we use the value obtained for *T* = 15 ∗ 65 bins (dashed red line)

**Extended Data Figure 11:**
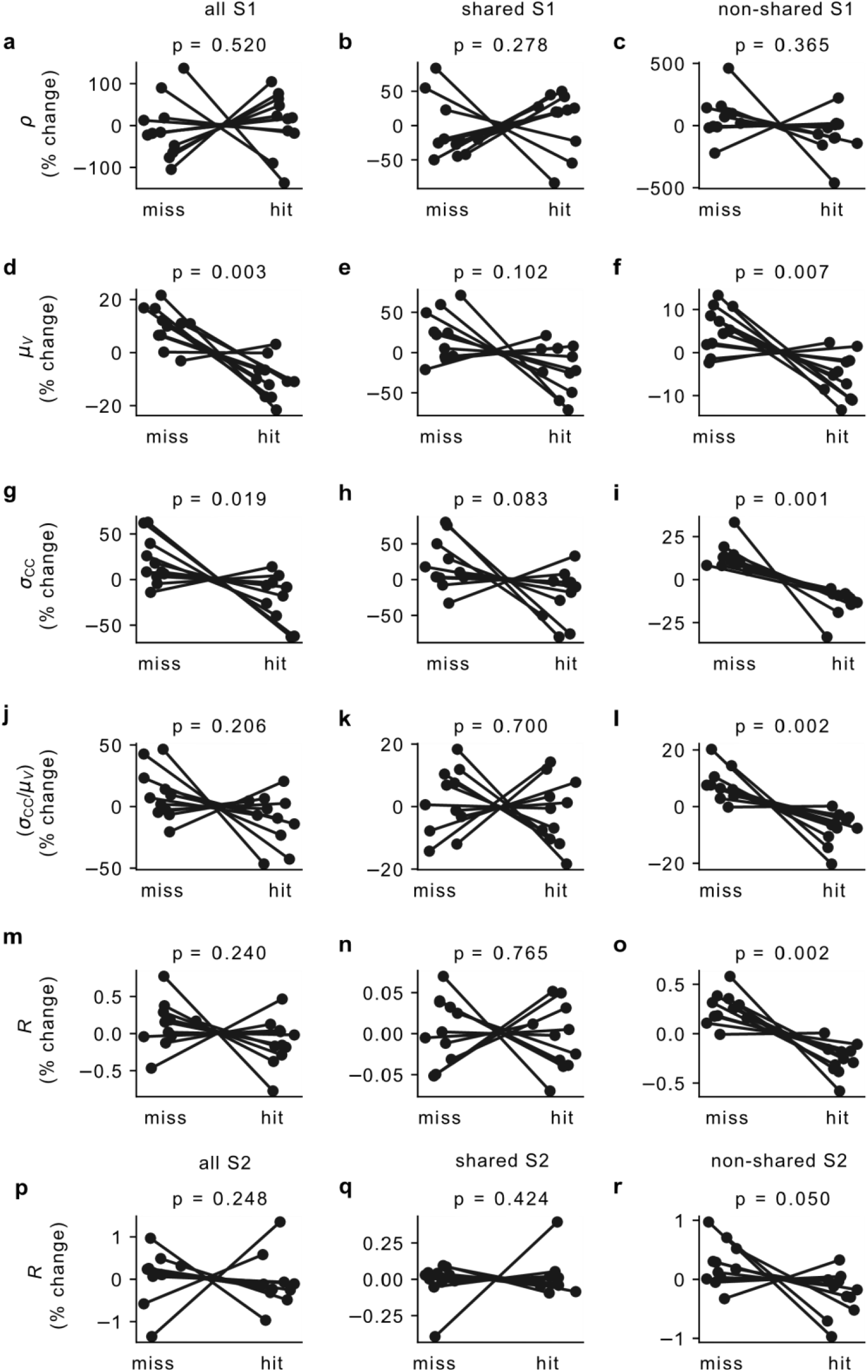
Recurrence analysis for all, shared and non-shared activity. The recurrence analysis results of **Fig. 5** are shown in more detail. The first row **(a-c)** shows the mean off-diagonal correlation *ρ*: for **(a)** all activity, **(b)** shared activity, and **(c)** non-shared activity. The second row **(d-f)** shows the mean on-diagonal covariance *µ*_*V*_. The third row **(g-i)** shows the standard deviation of the off-diagonal covariance *σ*_*CC*_. The fourth row **(j-l)** shows the ratio of the standard deviation of the off-diagonal covariance and the mean on-diagonal covariance *σ*_*CC*_*/µ*_*V*_. The fifth row **(m-o)** shows the change in recurrence *R*. The sixth row **(p-r)** also shows the change in the recurrence R, but estimated from neural activity in S2 instead of in S1.

**Extended Data Figure 12:**
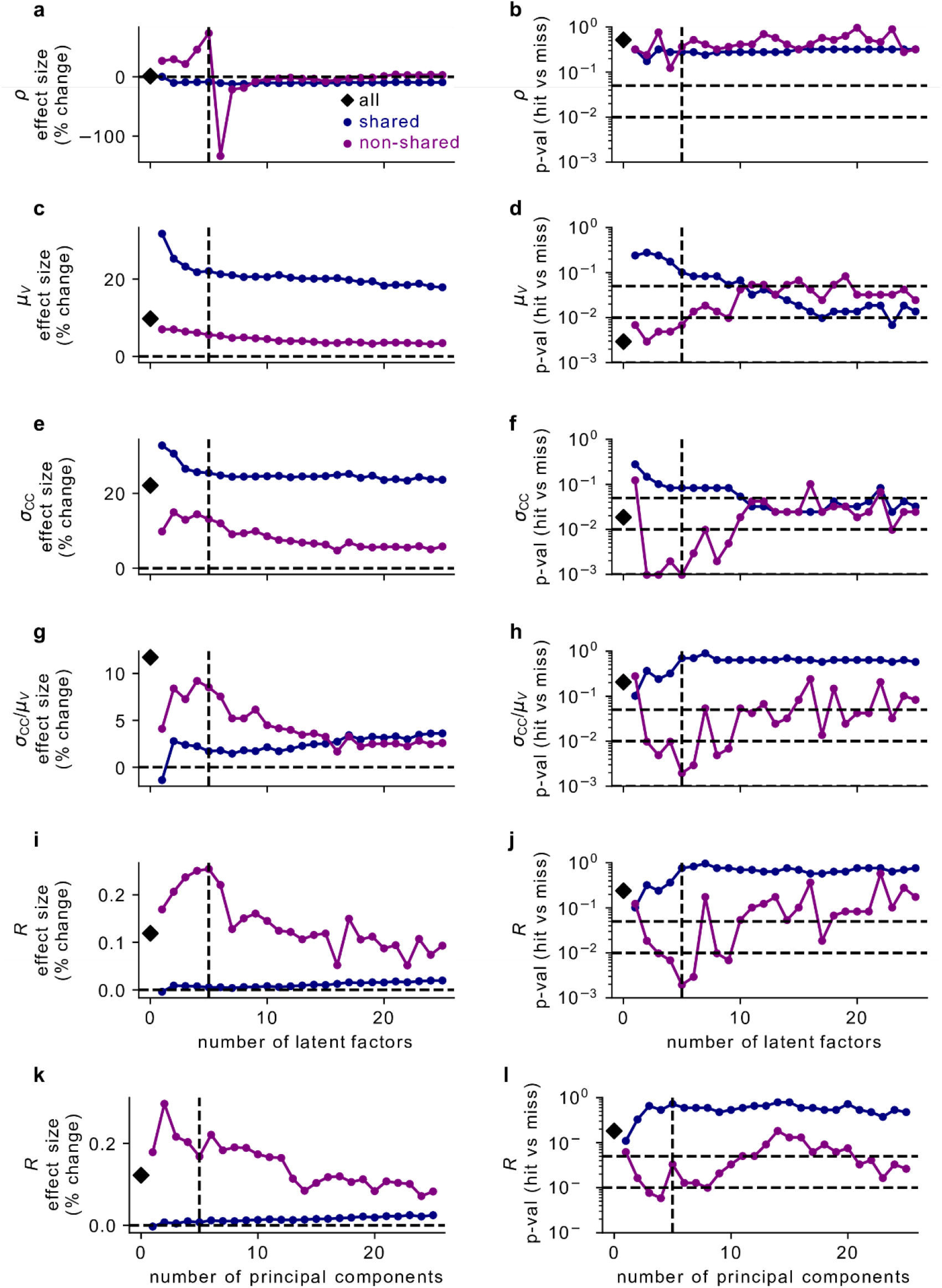
Recurrence analysis for different number of latent factors. The stability of the results shown in **Fig. 5** with respect to the choice of the number of latent factors is explored (dashed vertical line denotes the 5 latent factors we used in **Fig. 5**). The first row **(a-b)** explores the hit-miss difference for the mean off-diagonal correlation *ρ*; shown are **(a)** the effect size (dashed horizontal line denotes nil effect) and **(b)** the uncorrected p-value of the observed effect (dashed horizontal lines denote p=0.05, p=0.01, and p=0.001), for all activity (black diamond), shared activity (purple crosses) and non-shared activity (pink crosses). The second row **(c-d)** explores the hit-miss difference for the mean on-diagonal covariance *µ*_*V*_. The third row **(e-f)** explores the hit-miss difference for the standard deviation of the off-diagonal covariance *σ*_*CC*_. The fourth row **(g-h)** explores the hit-miss difference for the ratio of the standard deviation of the off-diagonal covariance and the mean on-diagonal covariance *σ*_*CC*_*/µ*_*V*_. The fourth row **(i-j)** explores the hit-miss difference for the recurrence *R*. The fifth row **(k-l)** shows the same hit-miss difference for R, but uses Principal Component Analysis (PCA) instead of LFA to estimate shared activity.

**Extended Data Figure 13:**
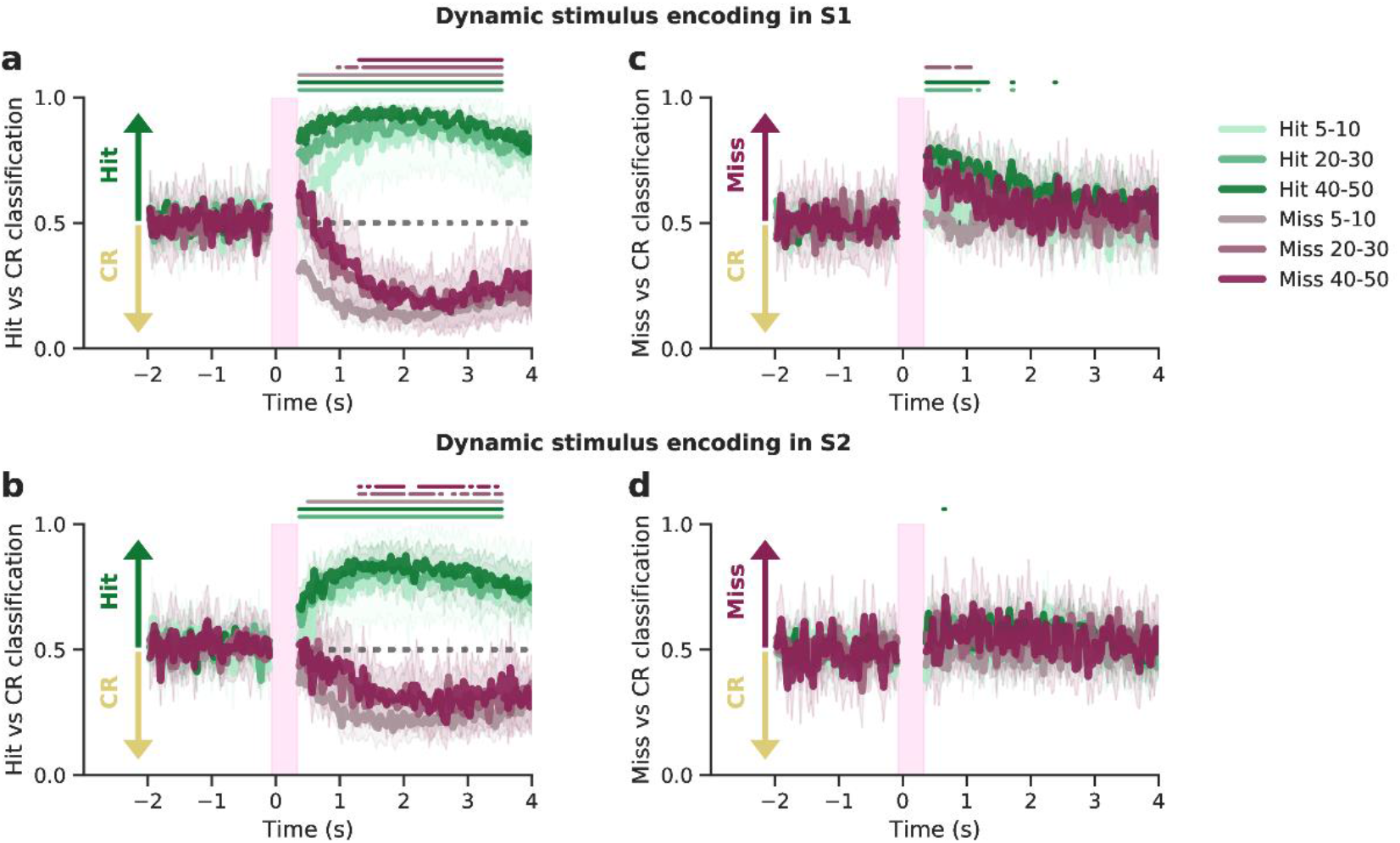
Dynamic decoding results split by number of cells targeted. The decoders of **Fig. 3** are shown, but additionally split by number of cells targeted. In other words, this figure shows the same data, but with an additional condition to separate trials by. Three ‘number of cells targeted’ conditions were used (instead of six) to increase data size, in particular for the rare scenarios (such as, e.g., ‘Miss 40-50’). Two recorded sessions did not include data for one trial type/cells targeted combination, and were therefore omitted (for that combination only).

**Extended Data Figure 14:**
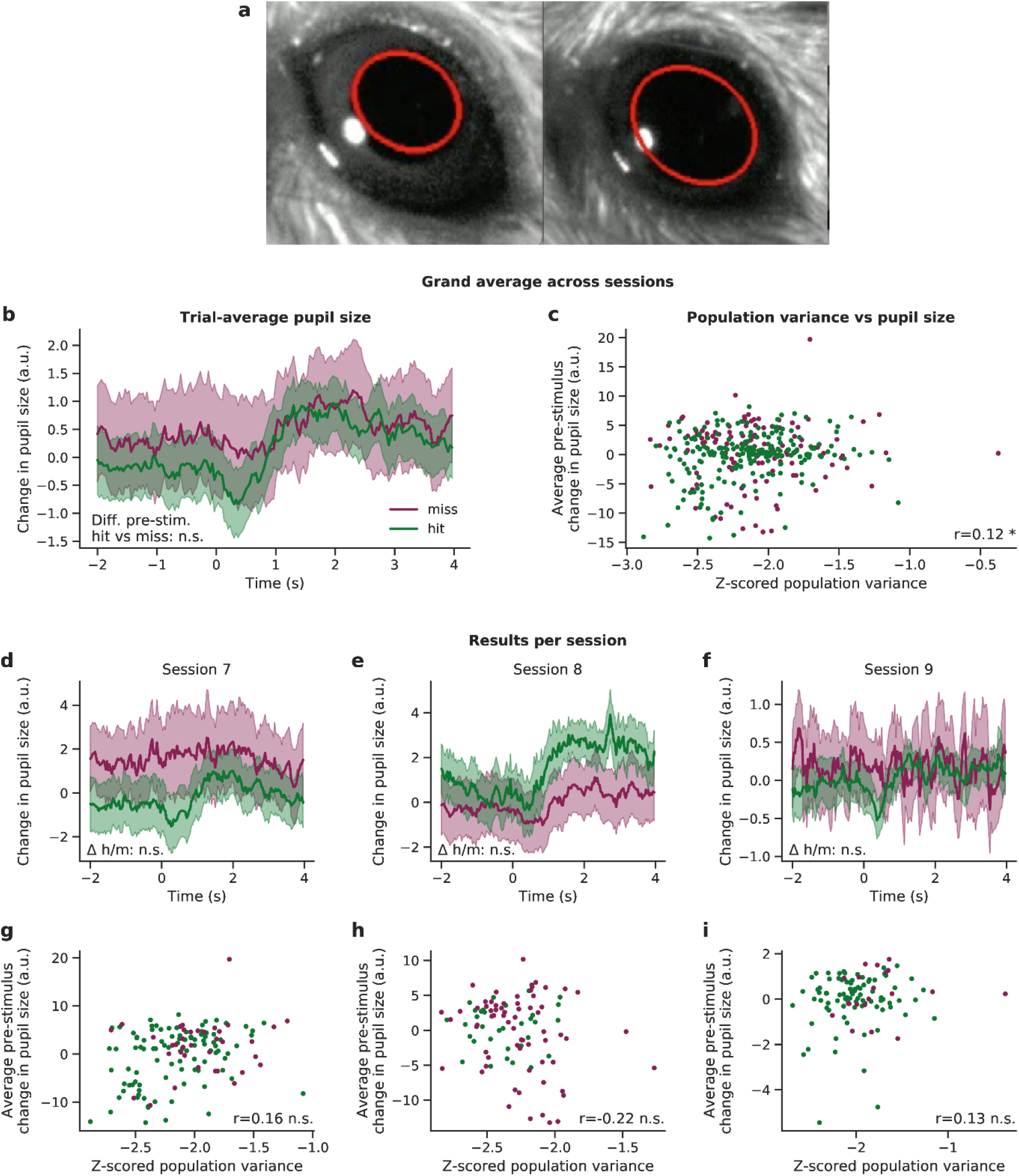
Pre-stimulus pupil size does not influence trial outcome. **(a)** Pupil size was measured for 3 sessions (sessions 7, 8 and 9, corresponding to Extended Data Fig. 4), see Methods. **(b)** Trial- and session-averaged pupil size dynamics for hit and miss trials, where shaded areas indicate the 95% confidence interval of the mean. The time-averaged pre-stimulus pupil size was not significantly different between hit and miss trials (two-sided t-test). **(c)** Population variance was very weakly correlated to pupil size across all sessions (both averaged across 500 ms pre-stimulus per trial) (Pearon correlation r=0.12, p=0.02). **(d-f)** Trial-averaged pupil size dynamics per session. Pre-stimulus differences between hit and miss were not significant for any of the sessions, although session 7 (panel d) almost reached the significance threshold (p=0.053). **(g-i)** Population variance vs pupil size per session. Pearson’ correlation was not significant for any of the sessions.

